# n-3 polyunsaturated fatty acids ameliorate post-infarction cardiac dysfunction through modulation of adiponectin-ceramide metabolism

**DOI:** 10.64898/2026.04.13.718333

**Authors:** Yuchen Liu, Wenqian Sun, Jia Liu, Hairui Wu, Peng Liu, Yanfang Chen, Run Zhang, Weirui Chen, Shuaishuai Wang, Xiaofei Guo, Wenzhong Zhang, Lihua Cao

## Abstract

**Background:** It has been shown that n-3 polyunsaturated fatty acids (n-3 PUFA) of marine origin exert significant beneficial effects on myocardial infarction (MI); however, the underlying mechanisms remained unclear. Ceramides play a vital role in the regulation of energy metabolism, mitochondrial function, and apoptosis. Through the integration of clincial studies and animal experiments, this study aimed to determine whether n-3 PUFA improved myocardial function by modulating ceramide metabolism.

**Methods:** In a case-control study, 100 patients with AMI and 100 healthy pariticipants were enrolled to measure serum ceramide concentrations. Meanwhile, mice were randomly allocated into 4 groups and administrated to a 3-week intervention with n-3 PUFA in triglyceride and phospholipid forms. A mouse model of MI was then established, followed by an additional 4 weeks of continuous intervention. Subsequent comprehensive assessments of cardiac function were performed in the mice. Finally, the mice were euthanized to conduct targeted ceramide lipidomic analysis and other relevant assays.

**Results:** The levels of serum C16:0-, C18:0-, C20:0-, C24:1-ceramides and total ceramides in patients with acute myocardial infarction (AMI) were significantly higher compared with the healthy controls. In the murine model of myocardial infarction, pathological analysis via TTC staining demonstrated that interventions with fish oil (triglyceride form) and krill oil (phospholipid form) both significantly reduced myocardial infarct size. Concomitant echocardiographic assessment confirmed that both treatments markedly elevated left ventricular ejection fraction (LVEF), with the magnitude of improvement being significantly superior to that of the model control group. Concurrently, compared with the model group, the concentrations of ceramides in cardiac tissue and serum were significantly lower in the groups with fish oil and krill oil intervention. Western blot analysis further confirmed that n-3 PUFA intervention upregulated adiponectin expression, reduced ceramide accumulation in myocardial tissue, and inhibited mitochondria-mediated cardiomyocyte apoptosis, thereby improving cardiac function and prognosis following myocardial infarction.

**Conclusions:** This work demonstrates that n-3 PUFA exert cardioprotective effects following MI mediated by adiponectin-ceramide axis. However, there is no significant difference regarding therapeutic efficacy of n-3 PUFA in triglyceride or phospholipid forms.

## 1. Introduction

Cardiovascular diseases (CVD) remain the leading cause of morbidity and mortality worldwide. It has been reported that CVD accounted for 2.2 million deaths among women and 1.9 million deaths among men across Europe in 2023^[1–2]^. Among CVD, acute coronary syndrome (ACS), typified by MI, often presents as the initial clinical manifestation. MI is an acute, life-threatening condition characterized by coronary artery occlusion, ischemia, and subsequent cell death in the affected cardiac region. As cardiomyocytes are permanent cells, cardiac tissue regeneration post-MI is severely limited, leading to impaired cardiac function and even heart failure (HF)^[3]^. Thus, there is an urgent need to identify safe and translatable strategies to improve structural and functional outcomes following MI.

Currently, to prevent MI and improve its prognosis, there are several therapeutic strategies to be proposed world wide such as inhibiting regulated cell death, attenuating inflammation, and inducing regeneration^[4,5]^. Concurrently, the n-3 poluunsaturated fatty acids (PUFA) of maine-origin to prevent and treat CVDs, including MI, has emerged as a prominent focus in cardiovascular research^[6,7]^. Epidemiological studies and randomized controlled trials have demonstrated that increased n-3 PUFA intake is associated with reduced MI-related mortality, with a negative correlation observed between intake levels and the risk of non-fatal MI and cardiac death^[8,9]^. While the underlying mechanisms remained complex and incompletely elucidated, key pathways include triglyceride reduction, inhibition of atherosclerotic plaque rupture, suppression of platelet aggregation, anti-inflammatory activity, and antioxidant effects^[10–14]^. Marine-derived n-3 polyunsaturated fatty acids (n-3 PUFAs), mainly including docosahexaenoic acid (DHA, 22:6n-3) and eicosapentaenoic acid (EPA, 20:5n-3), are primarily enriched in the human body in two natural forms: triacylglycerol (TAG) and phospholipid (PL), and are key nutritional components for maintaining a healthy human diet^[15]^. Although distinct n-3 PUFA forms may confer differential benefits in the management of metabolic diseases, whether they exert form-specific effects on MI outcomes remains unclear^[16]^.

Adiponectin is an endogenous bioactive protein derived from adipocytes, with diverse physiological functions such as promoting cardiomyocyte survival^[17]^. Accumulating evidence suggests that n-3 PUFA intervention can significantly elevate circulating adiponectin levels and regulate hepatic ceramide metabolism^[18]^. However, research on its direct effects on ceramide homeostasis in myocardial tissue and the underlying mechanisms remains relatively limited. Ceramide is formed via the dehydration of sphingosine and fatty acids and regulates cellular processes including differentiation, proliferation, apoptosis, senescence, and other vital biological activities^[19,20]^. As a bioactive lipid, ceramides are synthesized through three pathways: de novo synthesis, the sphingomyelinase pathway, and the salvage pathway^[21,22]^. In recent years, ceramides have garnered extensive attention across multiple research fields. Strong evidence links plasma ceramide levels to the development of coronary artery diseases^[23]^. Furthermore, substantial evidence suggests that ceramides are associated with vascular endothelial oxidative stress and dysfunction^[24–26]^, which are established independent predictors of poor cardiovascular prognosis. Overall, elevated plasma ceramide levels are generally detrimental to cardiovascular health. For example, plasma ceramide levels in patients during acute MI have been shown to predict the risk of complications, with concentrations correlating with a higher likelihood of MI recurrence and mortality. Recent studies have observed increased ceramide levels in cardiac tissue of rodents and humans during acute MI, and blocking de novo ceramide synthesis in rodents improves post-MI cardiac function^[27–30]^.

Previous studies have shown that n-3 PUFAs regulate ceramide metabolism via adiponectin^[18]^. However, the functional specificity and mechanisms of different n-3 PUFA forms within the adiponectin-ceramide axis remain unclear. To address this gap, we first conducted a case-control study. Non-targeted lipidomics revealed a marked elevation of C16:0 ceramide, which was confirmed by targeted ceramideomics: total serum ceramide and several individual species were significantly higher in cases than in controls. Based on these findings, we established a MI model in male C57BL/6J mice by left anterior descending (LAD) coronary artery ligation. This model was used to dissect how distinct n-3 PUFA forms differentially modulate ceramide profiles in cardiac tissue and circulation, and to determine their effects on post-MI cardiac function and prognosis.

## 2. Methods

### 2.1. Case–Control Study

#### 2.1.1 Study Design and Participants

For the case group, patients with AMI included in this study met the following criteria: 1) meeting the diagnostic criteria for ST-segment elevation myocardial infarction (STEMI) as specified in the 2025 American Heart Association (AHA) Guidelines^[31]^; 2) completion of admission assessment and blood sampling within 24 hours of symptom onset; 3) receipt of oral basic medications (e.g., aspirin) but no emergency percutaneous coronary intervention (PCI) or thrombolytic therapy; 4) no history of heart disease or any type of malignancy; 5) presence of severe, uncontrolled infection within the preceding 2 weeks (persistent body temperature > 38.5°C for > 48 hours). The healthy controls were randomly selected from individuals who were clinically healthy and had no prior history of cardiac disease. This study was approved by the Ethics Committee of the Medical College of Qingdao University (Registration Number: QDU-HEC-2024363).

#### 2.1.2 Clinical Characteristics and Serum Biochemistry

Clinical characteristics including gender, age, BMI, and blood pressure were recorded for patients upon admission. Following a 10-hour fasting, 10 mL of blood samples were collected into vacuum tubes the next morning and centrifuged at 3500 rpm for 10 minutes to isolate serum. Clinical parameters, namely triglycerides (TG), total cholesterol (TC), low-density lipoprotein cholesterol (LDL-C), high-density lipoprotein cholesterol (HDL-C), and fasting plasma glucose (FPG), were then quantified using an automatic biochemical analyzer, with results summarized in Table 1.

**Table 1.** Baseline Characteristics of the Study Population.

| Variables | Controls (n=100) | Cases (n=100) | P value |
| --- | --- | --- | --- |
| Male, n (%) | 58 (58) | 73(73) | 0.026 |
| Age | 58.9±14.1 | 61.5±12.5 | 0.109 |
| BMI, kg/m<sup>2</sup> | 25.66±4.32 | 25.14±3.05 | 0.351 |
| Smoke, n (%) | 12 (12) | 37 (37) | < 0.001 |
| Drink, n (%) | 10 (10) | 23 (23) | 0.013 |
| FHx, n (%) | 7 (7) | 16 (16) | 0.046 |
| Diabetes, n (%) | 40 (40) | 46 (46) | 0.391 |
| FPG, mmol/L | 6.03±2.03 | 6.77±3.11 | 0.084 |
| Systolic blood pressure, mmHg | 131±22 | 132±20 | 0.570 |
| Diastolic blood pressure, mmHg | 78±12 | 75±12 | 0.797 |
| TG, mmol/L | 1.48±0.89 | 1.56±0.98 | 0.479 |
| TC, mmol/L | 4.37±1.76 | 5.00±0.98 | 0.033 |
| HDL-C, mmol/L | 1.38±0.56 | 1.05±0.28 | 0.003 |
| LDL-C, mmol/L | 2.63±1.50 | 2.98±0.78 | 0.017 |

Characteristics of healthy control patients and acute myocardial infarction patients included in the study. Data are shown as mean±SD. Control vs AMI were compared using t tests for quantitative data and chi-square tests for categorical data. BMI indicates body mass index; FHx, family history; FPG, fasting plasma glucose; TG, triglyceride; TC, total cholesterol; HDL, high-density lipoprotein cholesterol; LDL, low-density lipoprotein cholesterol.

#### 2.1.3 Measurement of Cardiac Parameters by Echocardiography

Echocardiography was performed to assess the following clinical parameters: ejection fraction (EF), left ventricular end diastolic dimension (LVDd), interventricular septum diameter (IVS), left ventricular posterior wall (LVPW), left atrium anterior-posterior diameter (LAAP), and aortic valve annulus (AVA).

#### 2.1.4 Serum Lipidomic Analysis

Serum (20 μL) was mixed with 200 μL of dichloromethane-methanol (1:1, v/v), followed by vigorous vortexing for lipid extraction and homogenization. The mixture was centrifuged at 12,000 rpm for 15 min at 4 °C, and the resulting supernatant was transferred to a glass HPLC vial. The injection volume was set to 2 μL with a flow rate of 0.4 mL/min. Lipidomic metabolites were analyzed using an Agilent 6530C Q-TOF LC/MS system equipped with an ACQUITY UPLC BEH C18 column (2.1 mm × 100 mm, 1.7 μm). High-resolution mass spectrometry data were converted using ABF converter software. Metabolite identification was performed via Mass Spectrometry Data Independent Analysis (MS-DIAL) against a lipid database, and the results were integrated into the online platform (https://msflo.fiehnlab.ucdavis.edu). Principal component analysis (PCA) and volcano plot generation were conducted on the web-based platform https://www.metaboanalyst.ca/.

#### 2.1.5 Serum Targeted Ceramide Lipidomics Analysis

10 μL of serum was mixed with 100 μL of phosphate-buffered saline (PBS), followed by centrifugation at 12,000 rpm for 15 minutes at 4°C to collect the supernatant. A mixture of dichloromethane and methanol (1:1, v/v) was added to the supernatant at a volume ratio of 10:1, and the resulting solution was centrifuged at 12,000 rpm for 15 minutes at 4°C. To establish a standard curve, standard solutions of C16:0-, C18:0-, C20:0-, C22:0-, C24:0-, and C24:1-ceramides (ZZbio Co., Ltd., Shanghai, China) were serially diluted with dichloromethane-methanol (1:1, v/v) to generate gradient concentrations.

### 2.2 Animal Study

The animal study was approved by the Ethics Committee of the Medical College of Qingdao University (Registration Number: QDU-AEC-2024654).

#### 2.2.1 Experimental Design

Nine-week-old male C57BL/6J mice (Vital River Co., Ltd, Beijing, China) were housed under controlled conditions with a temperature range of 20-24 °C, relative humidity of 50–60%, and a 12-h light/dark cycle. All mice had ad libitum access to standard chow and sterile water. Following a 1-week acclimatization period, the mice were randomly assigned to 4 groups (n = 20 per group): (1) SHAM group: 0.1 mL of olive oil was administered via daily intragastric gavage; (2) MI group: 0.1 mL of olive oil was administered via daily intragastric gavage; (3) Krill oil group: 3 mL of a krill oil–water mixture (1:2, v/v) was administered via daily intragastric gavage; (4) Fish oil group: 0.1 mL of fish oil was administered via daily intragastric gavage. The fatty acid composition of the interventions is detailed in Supplementary Table 1.

After 3 weeks of intervention, surgical modeling was performed at week 4 using a small-animal respiratory anesthesia system (Shenzhen RWD Life Science & Technology Co., Ltd.). The procedure was adapted from the referenced literature and is briefly described as follows^[32]^: the sham-operated group underwent sham surgery (thoracotomy without coronary artery ligation), whereas the other three groups underwent permanent ligation of the left anterior descending coronary artery (LAD) at a site approximately 2 mm distal to the left atrial appendage via thoracotomy under blinded conditions to induce MI. Successful ligation was confirmed by immediate myocardial blanching; the thoracic cavity was then closed. Intervention was resumed at week 5 and continued for an additional 4 weeks. Cardiac function was serially assessed by transthoracic echocardiography throughout this period.

At the end of the experimental period, mice were fasted overnight and euthanized. Heart tissues were harvested immediately: one portion was snap-frozen in liquid nitrogen and stored at −80 °C for biochemical analyses, while the other portion was fixed in 4% paraformaldehyde for subsequent histological examinations. The experimental design is illustrated in Figure 1.

**Figure 1.**
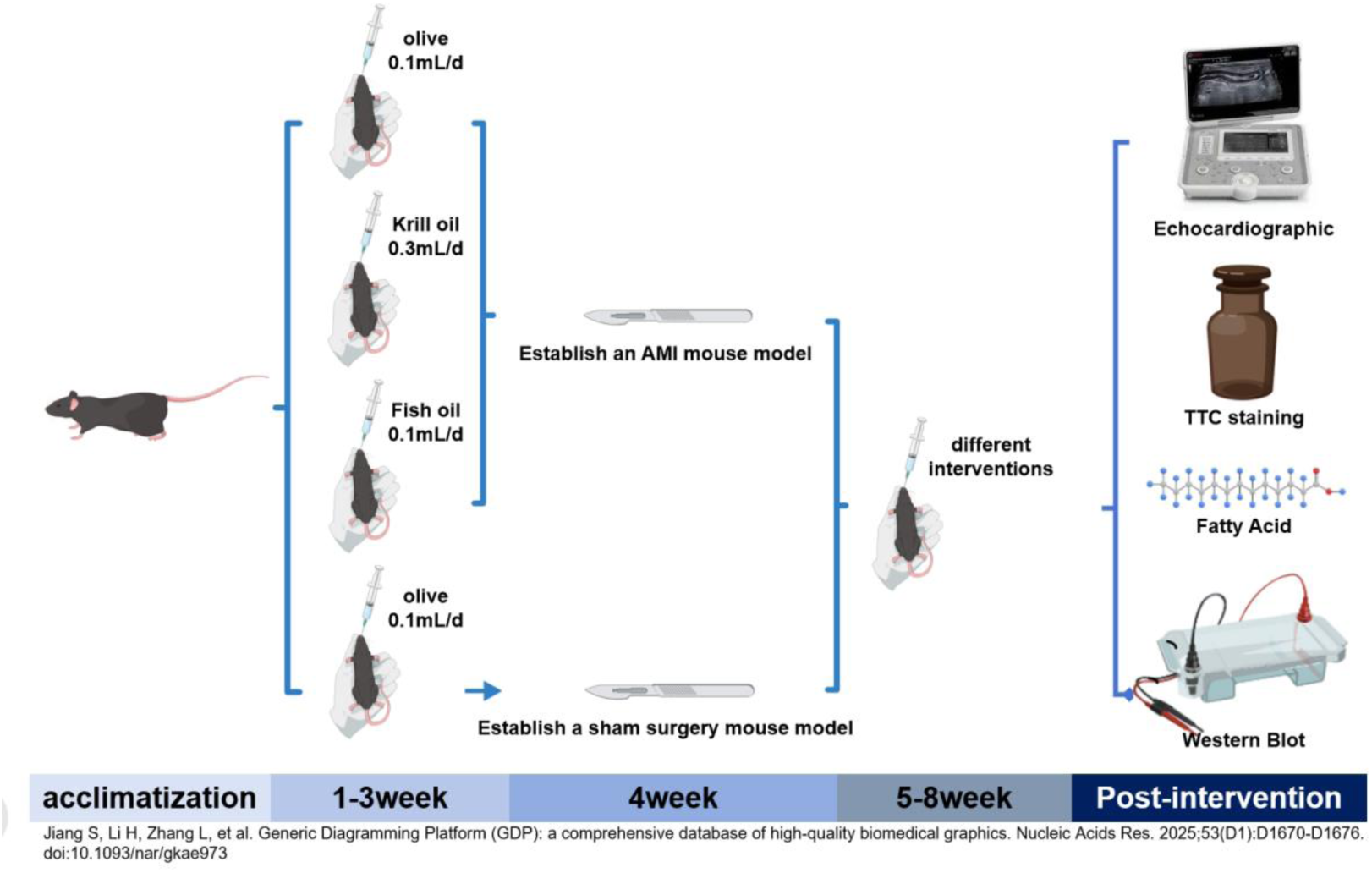
Overview of the animal experiment flow chart.

#### 2.2.2 Echocardiographic Measurements

Mice were anesthetized by intraperitoneal injection of 1% pentobarbital sodium. Subsequently, two trained ultrasound operators independently performed transthoracic echocardiography under blinded conditions using a small animal ultrasound imaging system (VINNO 6 LAB, Suzhou) equipped with a 23 MHz linear probe. The following echocardiographic parameters were detected, including ejection fraction; interventricular septum diameter at diastole (IVSd); left ventricular internal dimension at diastolic end (LVIDd); LV posterior wall thickness at diastole (LVPWd); interventricular septum diameter at systole (IVSs); left ventricular internal dimension at systolic end (LVIDs); LV posterior wall thickness at systole (LVPWs); heart rate; Left Ventricular End-Diastolic Volume (LVEDV); Left Ventricular End-Systolic Volume (LVESV); stroke volume; cardiac output and fractional shortening.

#### 2.2.3 Determination of Myocardial Infarction Size

The myocardium was sectioned into 1.0 mm-thick transverse slices from the apex to the cardiac base, followed by staining with triphenyltetrazolium chloride (TTC) solution at 37 ℃ for 30 mins. After thorough rinsing with PBS buffer, the slices were visualized under a Nikon Eclipse E400 phase-contrast microscope (Nikon Corporation, Tokyo, Japan). Viable myocardium exhibited a brick-red staining, whereas infarcted regions remained unstained (appearing gray). The percentage of myocardial infarct area was quantitatively determined using ImageJ software (version 1.8.0).

#### 2.2.4 Cardiac Fatty Acid Composition Analysis

Briefly, 0.5 mL of serum was subjected to lipid extraction with methanol/chloroform (1:1, v/v) at 4 ℃ for 24 h. Following esterification with methanol/sulfuric acid (0.9 mol/L) and toluene at 70 ℃, fatty acid methyl esters were quantified using an Agilent 7820A gas chromatograph equipped with an Agilent DB-23 capillary column (60 m, 0.25 mm × 0.25 μm)^[33]^. Cardiac fatty acid composition is presented in Supplementary Table 2.

#### 2.2.5 Cardiac Lipidomic Analysis

Cardiac tissue (∼ 10 mg) was homogenized in 100 μL PBS and centrifuged at 12,000 rpm for 15 min at 4 °C to collect the supernatant. The supernatant (100 μL) was mixed with 1000 μL of dichloromethane–methanol (1:1, v/v), followed by vortexing and centrifugation at 12,000 × g for 15 min at 4 °C. The lower organic phase was transferred to a glass HPLC vial. All other experimental conditions followed those described in Section 2.1.4.

#### 2.2.6 Targeted Ceramide Lipidomics Analysis

Briefly, 20 mg of heart tissue was homogenized with 200 µL of PBS twice at 60 Hz for 60 seconds per cycle. The homogenate was centrifuged at 12,000 rpm for 15 minutes at 4 ℃ to collect the supernatant. Subsequently, 100 µL of the supernatant was mixed with 1000 µL of dichloromethane/methanol (1:1, v/v), vortexed for 30 seconds, and centrifuged again at 12,000 rpm for 15 minutes. The injection volume was set to 2 µL with a flow rate of 0.4 mL/min. Other experimental conditions were consistent with those described in Section 2.1.5.

#### 2.2.7 Western Blot assay

Approximate 50 mg heart tissue was homogenized in 250 μL utilizing lysate solution (Shanghai Epizyme Biomedical Technology Co., Ltd, China) and subsequently centrifuged at 12,000 rpm for 10 minutes at 4 ℃. Protein concentration was determined via the bovine serum albumin (BSA) assay. Proteins were separated by sodium dodecyl sulfate-polyacrylamide gel electrophoresis (SDS-PAGE, 8-10%) and transferred onto polyvinylidene fluoride (PVDF) membranes (Millipore, Darmstadt, Germany). Following blocking with 5% non-fat milk, the membranes were incubated overnight at 4 °C with primary antibodies, including adiponectin (1:1000 dilution, Affinity Biosciences Co, Jiangsu), adiponectin receptor 1 (AdipoR1) (1:1000 dilution, Affinity Biosciences Co, Jiangsu), adiponectin receptor 2 (AdipoR2) (1:1000 dilution, Affinity Biosciences Co, Jiangsu), SPTLC2 (1:1000 dilution, Santa Cruz Biotechnology (Shanghai) Co., Ltd.), SPTLC3 (1:1000 dilution, ZENBIO Co., Chengdu), SMPD3 (1:1000 dilution, ZENBIO Co., Chengdu), ceramide synthase 6 (1:1000 dilution, ZENBIO Co., Chengdu), acid ceramidase (ASAH1) (1:1000 dilution, ZENBIO Co., Chengdu), n-acylsphingosine amidohydrolase 3 (ASAH3) (1:2000 dilution, Affinity Biosciences Co, Jiangsu), n-acylsphingosine amidohydrolase 3-like (ASAH3L) (1:1000 dilution, Affinity Biosciences Co, Jiangsu), alkaline ceramidase 3 (PHCA) (1:1000 dilution, Affinity Biosciences Co, Jiangsu), mitochondrial fission factor (MFF) (1:1000 dilution, Abcam Co., USA), mitofusin 1 (Mfn1) (1:1000 dilution, Abcam Co., USA), mitofusin 2 (Mfn2) (1:1000 dilution, Abcam Co., USA), optic atrophy 1 (OPA1) (1:1000 dilution, Abcam Co., USA), recombinant tumor protein p53 (p53) (1:1000 dilution, Affinity Biosciences Co, Jiangsu), caspase 3 (1:1000 dilution, Affinity Biosciences Co, Jiangsu), cleaved caspase-3 (1:1000 dilution, Affinity Biosciences Co, Jiangsu), B-cell lymphoma 2 (Bcl-2) (1:1000 dilution, Affinity Biosciences Co, Jiangsu), Bcl-2 associated X protein (BAX) (1:2000 dilution, Affinity Biosciences Co, Jiangsu), and glyceraldehyde-3-phosphate dehydrogenase (GAPDH) (1:5000 dilution, Affinity Biosciences Co, Jiangsu). Membranes were washed three times with Tris-buffered saline (TBS) containing 0.05% Tween-20 (TBST) and then incubated with secondary antibodies for 1.5 hours. Immunoreactive signals for protein expression were detected using a chemiluminescence kit on an image analyzer (Tanon-5200, Tanon Science & Technology Co., Ltd, Shanghai, China).

#### 2.2.8 Statistical Analysis

Statistical analyses were performed as follows: normality of data distribution was assessed using the Shapiro–Wilk test. For data that did not meet the assumption of normality, non-parametric tests were applied—namely, the Mann–Whitney U test for two-group comparisons and the Kruskal–Wallis test with Dunn’s post hoc correction for multiple groups. For ANOVA, if Levene’s test indicated a violation of homogeneity of variances, Welch’s ANOVA followed by the Games–Howell post hoc test was used. Differences between two groups of quantitative data were evaluated using the unpaired Student’s t-test; comparisons among three or more groups were conducted using one-way analysis of variance (ANOVA), followed by Bonferroni’s post hoc test for multiple comparisons, assuming homogeneity of variances (assessed by Levene’s test). Categorical variables were analyzed using the chi-square test (χ² test) in SPSS 26.0. Correlation analyses between cardiac functional parameters and serum ceramide concentrations, Kaplan–Meier survival curves, and all graphical representations were generated using GraphPad Prism 9.5.0. Data are presented as mean ± standard deviation (mean ± SD). A two-tailed P-value < 0.05 was considered statistically significant.

## 3. Results

### 3.1 Echocardiographic Parameters of Cases and Controls

Echocardiographic measurements showed that interventricular septum diameter (IVS), left atrium anterior-posterior diameter (LAAP), and aortic valve annulus (AVA) were significantly higher in patients with MI compared with the healthy controls (Figures 2C, E, and F). More importantly, ejection fraction (EF) was significantly lower in the cases than in the controls (Figure 2A). Besides, no statistically significant difference was observed between the two groups regarding left ventricular end-diastolic diameter (LVDd) and left ventricular posterior wall thickness (LVPW) (Figures 2B and 2D).

**Figure 2.**
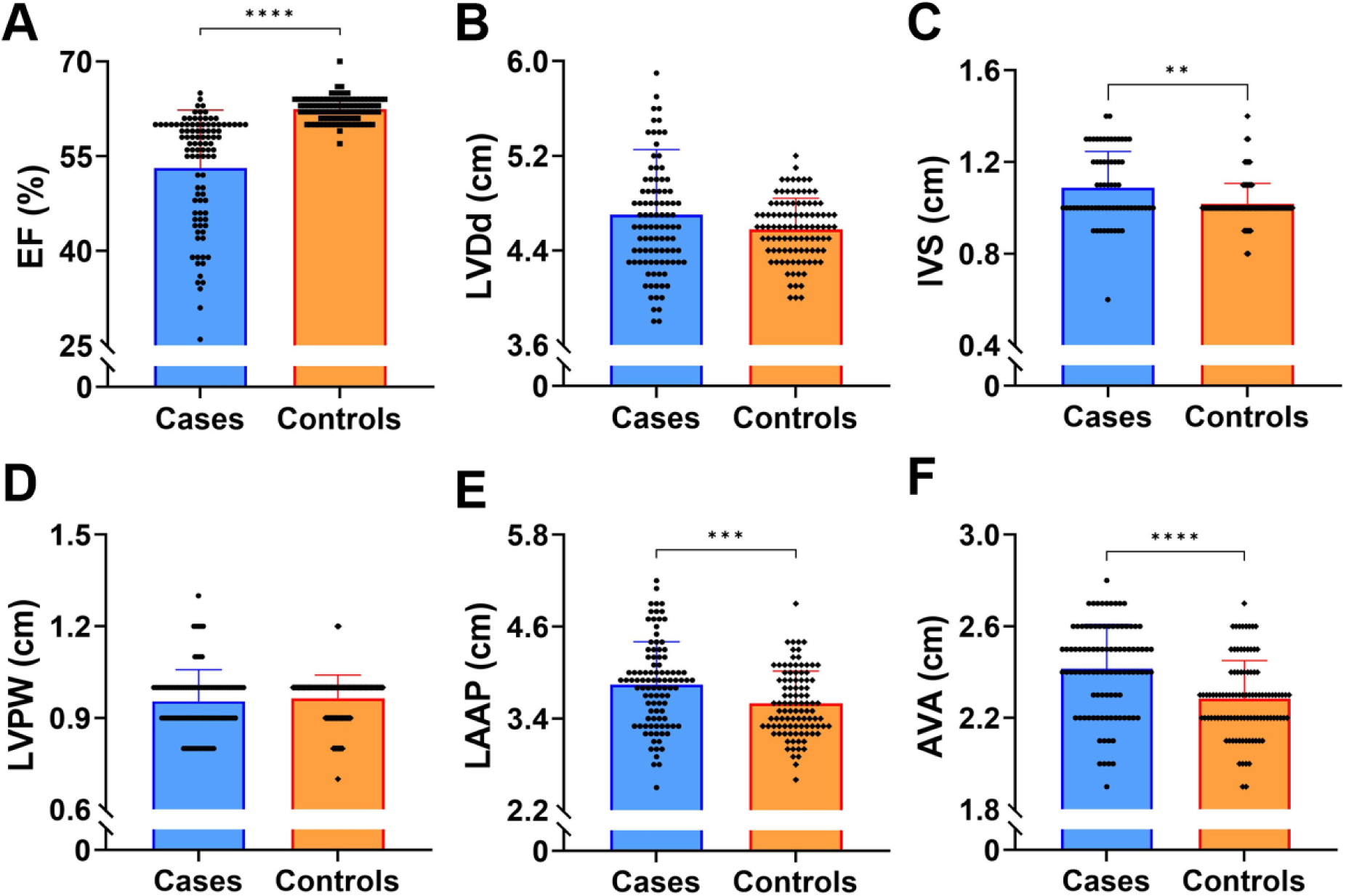
Echocardiographic parameters of patients with controls and AMI. (A) ejection fraction (EF). (B) left ventricular end diastolic dimension (LVDd). (C) interventricular septum diameter (IVS). (D) left ventricular posterior wall (LVPW). (E) left atrium anterior-posterior diameter (LAAP). (F) aortic valve annulus (AVA). Data are presented as the mean ± SD. **P<0.01, ***P<0.001, ****P<0.0001 by t test.

### 3.2 Correlation Analysis of Serum Ceramide Concentrations Between Case and Control Groups

In PLS-DA, the case and control groups were largely distinguishable, with the PC1 and PC2 explaining 65.4% and 13.2% of the variance, respectively (Figure 3A). A volcano plot, which visualizes log2 fold changes against -log10 p-values, revealed that serum C16:0-ceramide levels were significantly higher in the case group compared to the control group (Figure 3B). Targeted ceramide profiling showed that serum levels of C16:0-, C18:0-, C20:0-, C24:1-, and total ceramides were significantly higher in the cases than in the controls, whereas C22:0- and C24:0-ceramide levels were significantly reduced (Figure 3). Correlation analysis demonstrated that cardiac troponin I (cTnI) was positively correlated with total serum ceramide concentrations (Figure 4A). Ejection fraction (EF) showed negative correlations with serum C16:0-, C18:0-, C20:0-, C24:1-, and total ceramide concentrations (Figures 4B-E, and H), while exhibiting positive correlations with C22:0- and C24:0-ceramides (Figures 4F and G).

**Figure 3.**
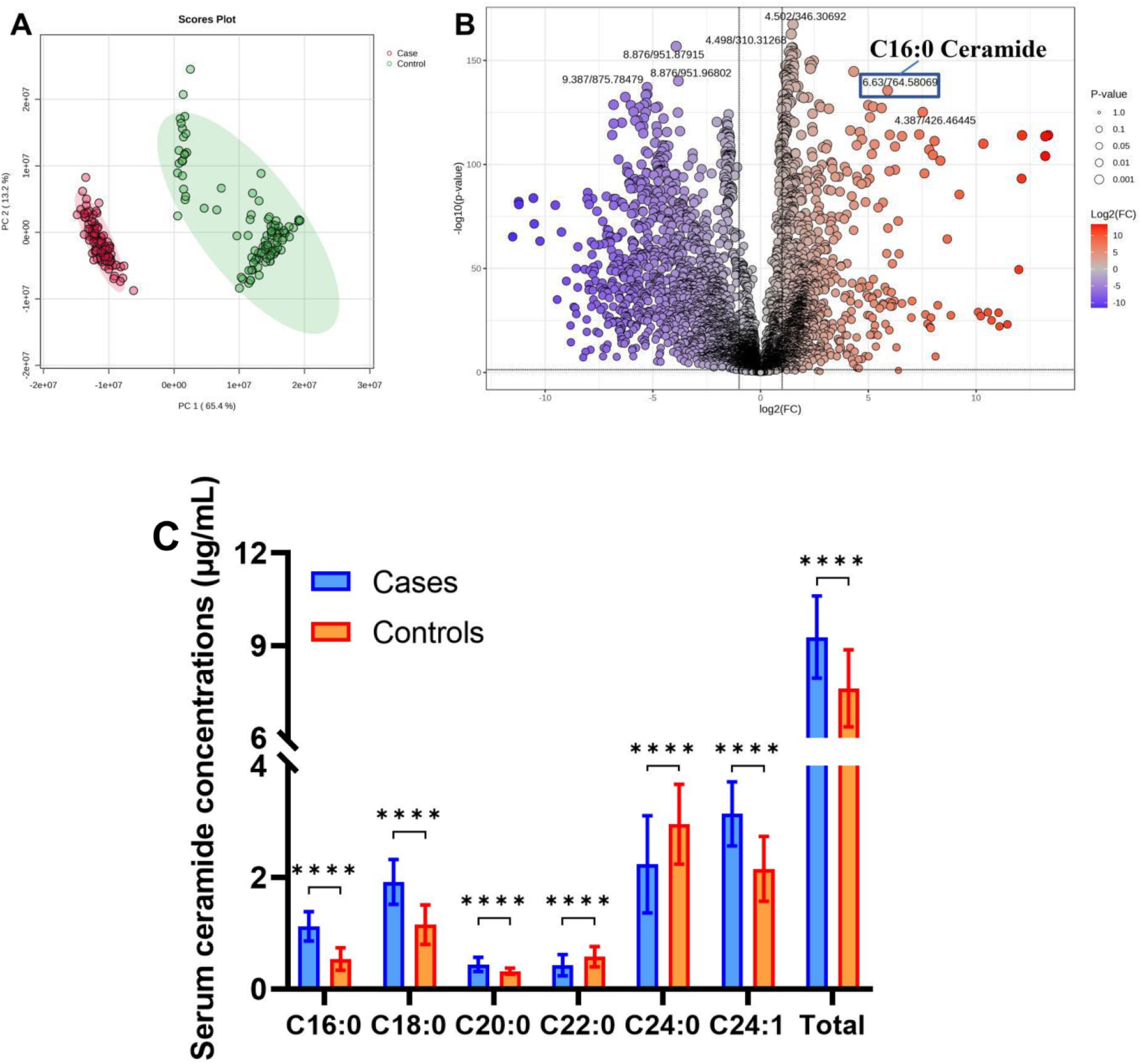
Serum lipids were analyzed using lipidomics and targeted ceramide lipidomics approaches. (A) PCA of serum lipids between the case and control groups. (B) Volcano plot of serum lipid metabolic data comparing the case and control groups. (C) Ceramide concentrations in control and case patients. Data are presented as the mean ± SD. ****P<0.0001 by t test.

**Figure 4.**
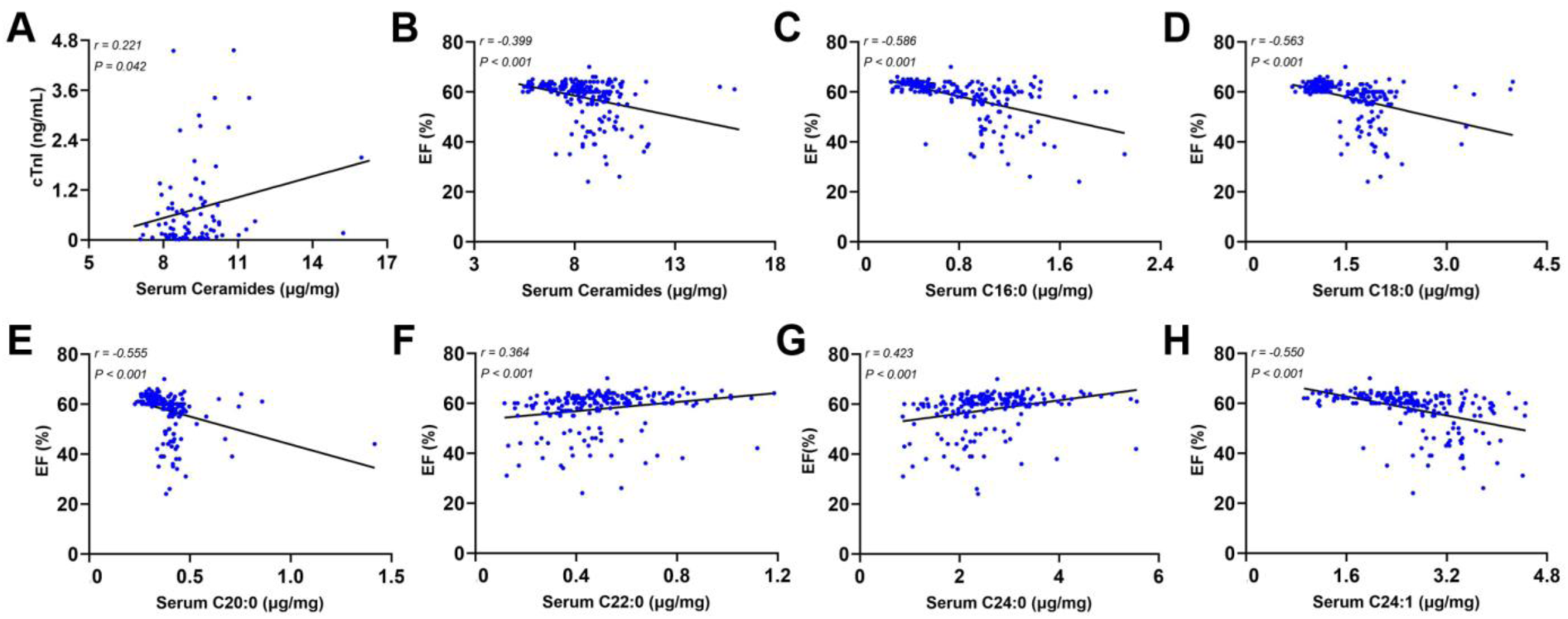
(A) Correlative analysis of cTnI with Serum Ceramides. (B-H) Correlative analysis of ejection fraction (EF) with Serum Ceramides (B), Serum C16:0 (C), Serum C18:0 (D), Serum C20:0 (E), and Serum C22:0 (F), Serum C24:0 (G), Serum C24:1 (H). Correlations were assessed by parametric Pearson’s test.

### 3.3 Administration of fish oil and krill oil improved myocardial infarction prognosis in mice

With the exception of one mouse in the sham-operated group that died due to gavage error, all other mortalities occurred during the perioperative period (i.e., from the initiation of surgery to 6 hours postoperatively)^[32]^ and the acute phase of myocardial infarction (within 2 weeks). Specifically, during the perioperative period, 3 mice in the SHAM group, 7 in the MI group, and 5 in each of the krill oil and fish oil groups succumbed. At the conclusion of the intervention, the mortality rate in the myocardial infarction group was significantly higher than that the sham-operated group, whereas the mortality rates in the krill oil and fish oil groups were lower than that in the myocardial infarction group (Figure 5). Furthermore, TTC staining showed that myocardial infarct size was significantly greater in the MI group compared with the SHAM group. However, adminstration of fish oil and krill oil significantly reduced infarct size in comparison with the MI group (Figure 6).

**Figure 5.**
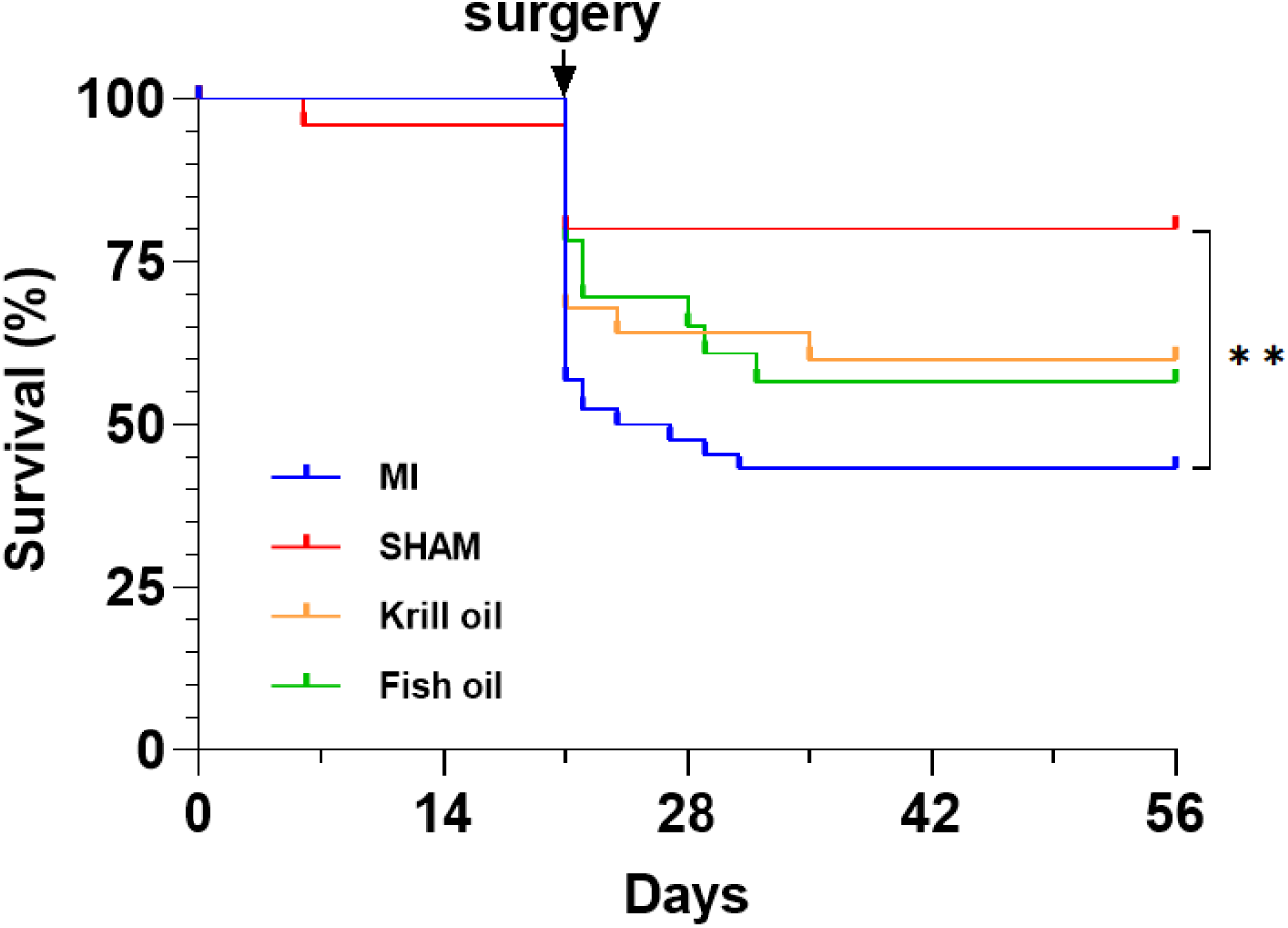
Kaplan-Meier survival curve for C57BL/6J mice subjected to MI or sham surgery and treated with olive, fish or krill oil. **P<0.01 MI vs sham by pairwise log-rank test.

**Figure 6.**
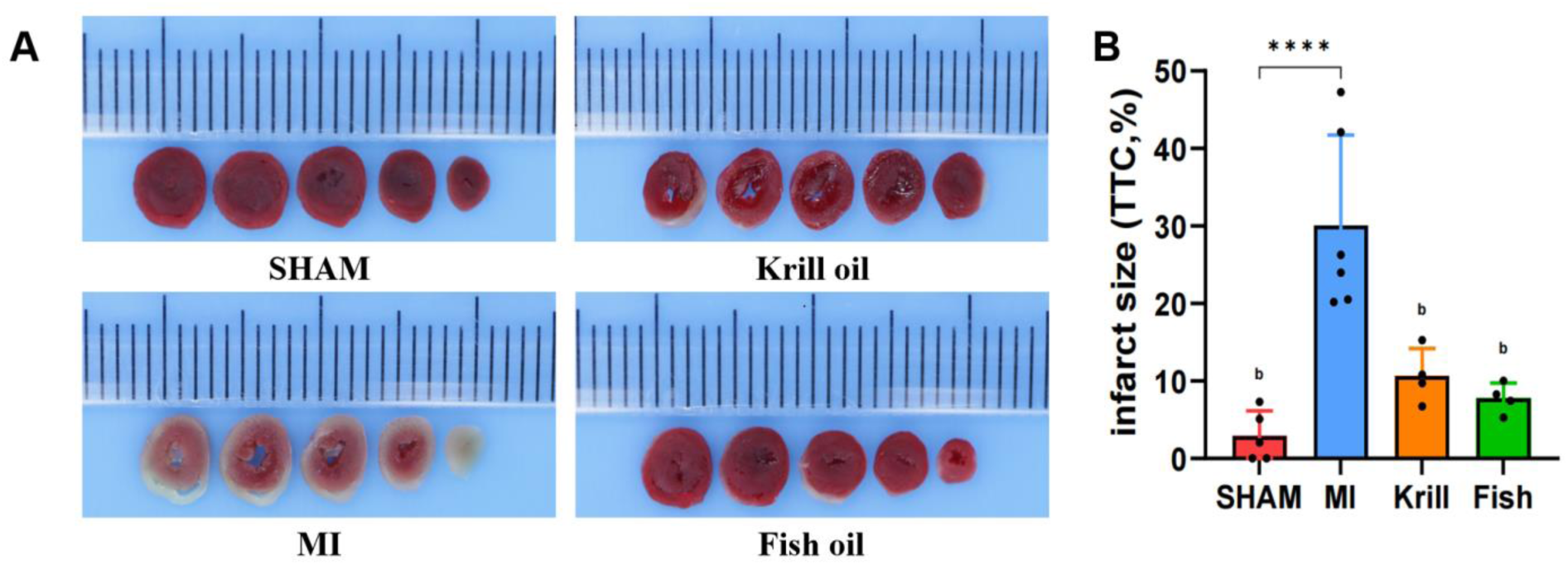
Images of tetrazolium chloride (TTC) stained heart sections (A) and quantification (B) of the 4 groups. Data are presented as the mean ± SD. Different letters indicate significant difference between groups.

### 3.4 Administration of Krill oil and fish oil improved cardiac function in mice

Echocardiographic analysis showed that the MI group exhibited significantly reduced ejection fraction, fractional shortening, IVSs, and LVPWs, concomitant with significantly increased LVIDs and LVESV compared with the SHAM group (Fig. 7B, F, G, H, J, M). However, administration of krill oil and fish oil significantly ameliorated MI-induced cardiac dysfunction, as evidenced by restored ejection fraction and fractional shortening (Fig. 7B, M) and normalized LVIDs and LVESV (Fig. 7G, J). In contrast, no significant difference was observed between groups regarding IVSd, LVIDd, LVPWd, stroke volume, or cardiac output (Fig. 7C, D, E, K, L).

**Figure 7.**
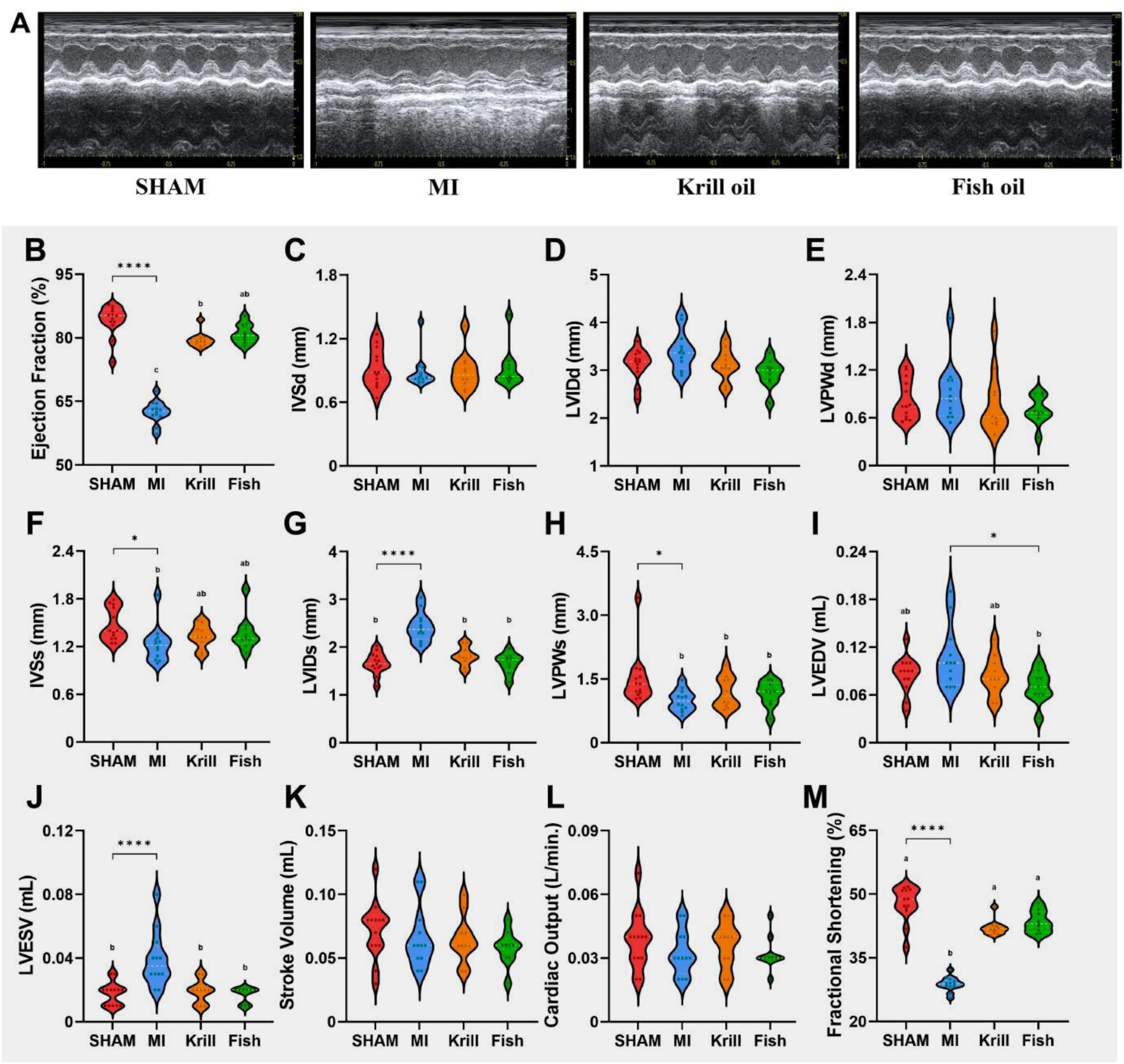
Effect of krill and fish oil intervention on cardiac function in C57BL/6J mice (A). Representative left ventricular M-mode echocardiographic images. (B) ejection fraction; (C) interventricular septum diameter at diastole (IVSd); (D) left ventricular internal dimension at diastolic end (LVIDd); (E) LV posterior wall thickness at diastole (LVPWd); (F) interventricular septum diameter at systole (IVSs); (G) left ventricular internal dimension at systolic end (LVIDs); (H) LV posterior wall thickness at systole (LVPWs); (I) Left Ventricular End-Diastolic Volume (LVEDV); (J) Left Ventricular End-Systolic Volume (LVESV); (K) stroke volume; (L) cardiac output and (M) fractional shortening. Data are presented as the mean ± SD. Different letters indicate significant difference between groups.

### 3.5 Effects of Krill Oil and Fish Oil on Lipid Metabolism and Cardiac Ceramide Profiles

PLS-DA revealed a distinct separation trend in metabolic profiles of myocardial tissue and serum samples between the MI group and the SHAM group (Supplementary Figure 1A, C). Screening of differential metabolites demonstrated that levels of C16:0-ceramide and C18:0-ceramide in both myocardial tissue and serum were significantly elevated in the MI group compared to the SHAM group (Supplementary Figure 1B, D). Targeted ceramide lipidomics further confirmed that myocardial tissue levels of C16:0-, C18:0-, C24:0-, C24:1-ceramides, as well as total ceramides, were significantly higher in the MI group relative to the SHAM group. Notably, intervention with krill oil or fish oil resulted in a significant reduction in myocardial C16:0-, C18:0-ceramides, and total ceramides. In serum, all detected ceramide subtypes—with the exception of C22:0-ceramide—were significantly increased in the MI group compared to the SHAM group (Figure 8A, B). Following intervention, serum levels of all ceramide subtypes (excluding C16:0-ceramide) and total ceramides were significantly decreased relative to the MI group (Figure 8C, D).

**Figure 8.**
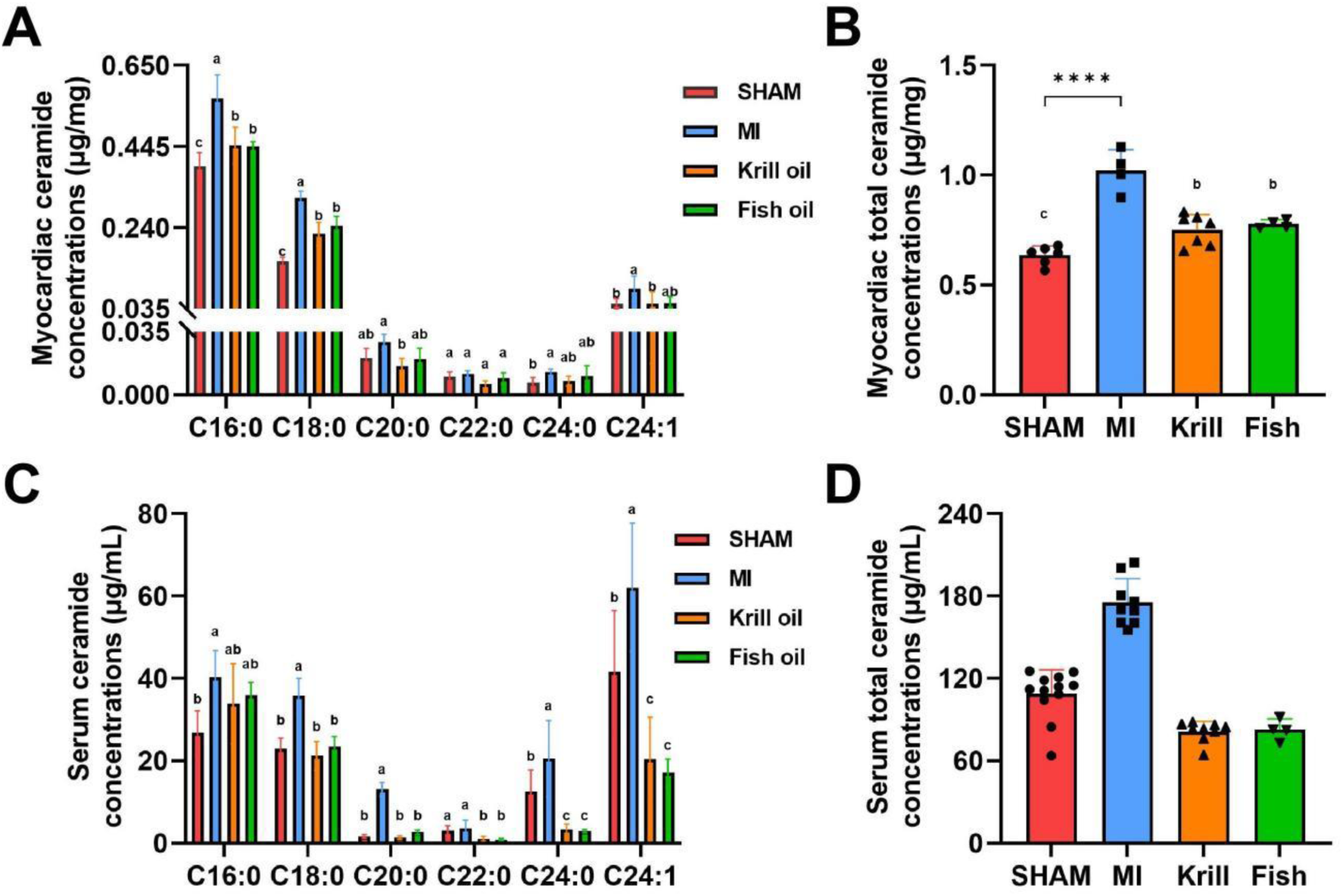
Effect of krill and fish oil administration on myocardial (A, B) and serum (C, D) ceramide concentrations in male C57BL/6J mice. Data are presented as the mean ± SD. Different letters indicate significant difference between groups.

### 3.6 Administration of Krill Oil and Fish Oil on Protein Expression Levels

Compared with the SHAM group, the protein expression levels of adiponectin and AdipoR2 were significantly reduced in the cardiac tissues of mice in the MI group. Notably, administration of krill oil and fish oil markedly reversed this downregulation, leading to significantly elevated protein levels of both targets relative to the MI group (Figure 9A). Correspondingly, regarding the ceramide metabolic pathway, the cardiac protein expression levels of SPTLC2 and SPTLC3 in the krill oil and fish oil groups were significantly lower compared with the MI group, whereas the opposite pattern was observed for the catabolic enzyme ASAH1 (Figures 9B, 9C). For mitochondrial dynamics-related proteins, the protein expression levels of MFF were significantly lower in the krill oil and fish oil groups compared with the MI group (Figure 9D). At the level of apoptosis regulation, krill oil and fish oil intervention significantly downregulated the protein expression of pro-apoptotic proteins p53 and caspase-3, with statistically significant differences compared to the MI group (Figure 9E).

**Figure 9.**
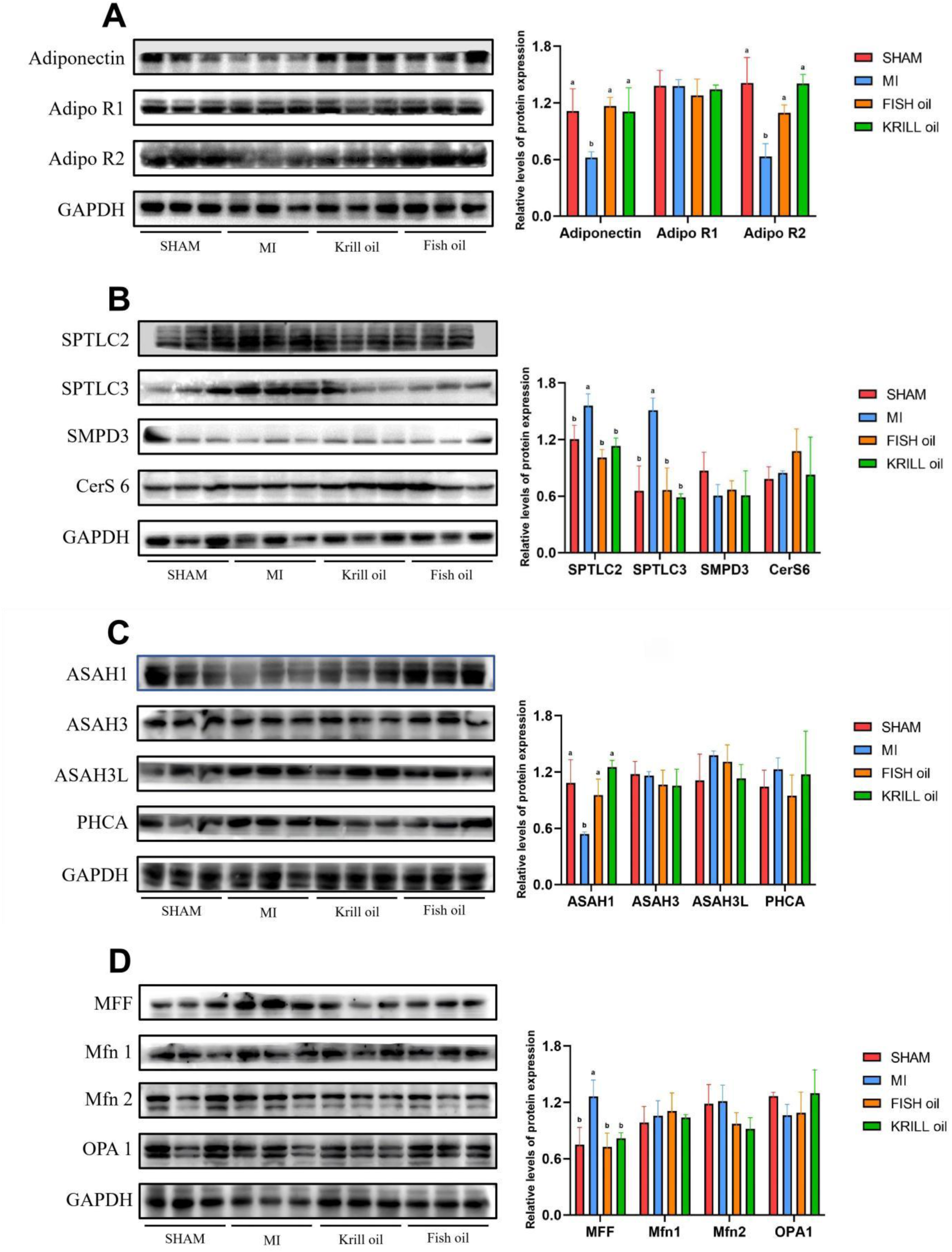
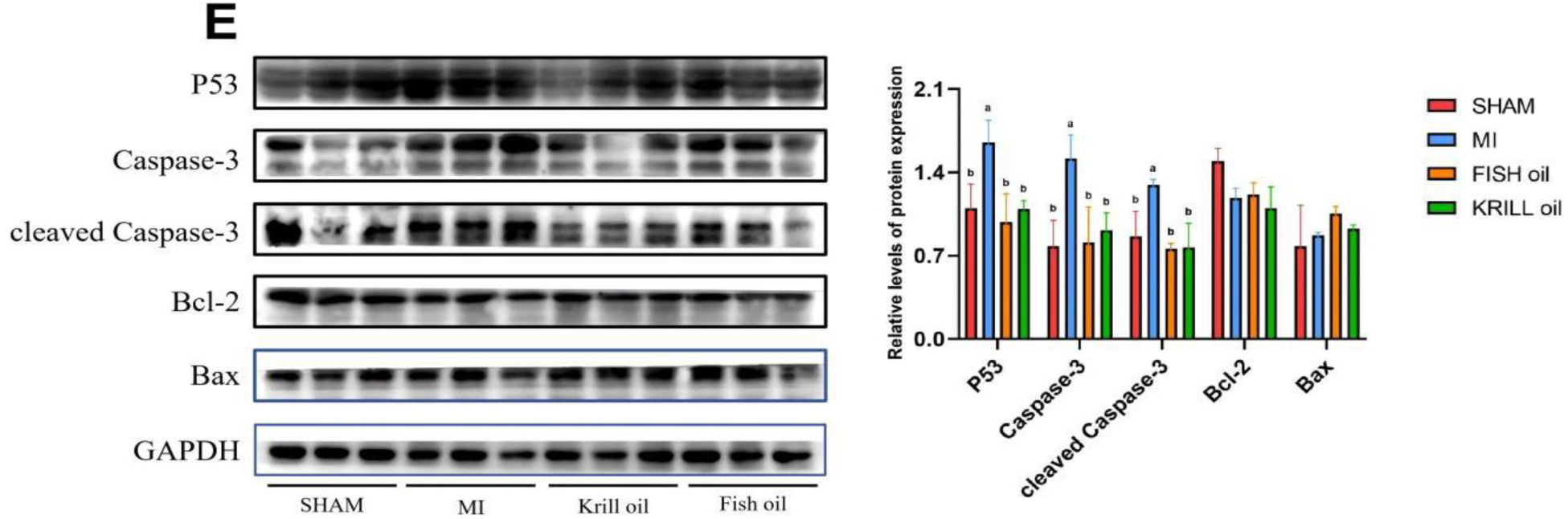
Effect and quantification of n-3 PUFA treatment on cardiac protein expression levels. (A) Adiponectin, AdipoR1, AdipoR2. (B) SPTLC2, SPTLC3, SMPD3, CerS6. (C) ASAH1, ASAH3, ASAH3L, PHCA. (D) MFF, Mfn1, Mfn2, OPA1. (E) P53, Caspase-3, cleaved Caspase-3, Bcl-2, Bax, expression. Data are presented as the mean ± SD. Different letters indicate significant difference between groups. Abbreviations: AdipoR1: adiponectin receptor protein 1; AdipoR2: adiponectin receptor protein 2; SPTLC2: Serine Palmitoyltransferase Long Chain Base Subunit 2; SPTLC3: Serine Palmitoyltransferase Long Chain Base Subunit 3; SMPD3: Sphingomyelin phosphodiesterase 3; CerS6: Ceramide Synthase 6; ASAH1: Acid ceramidase; ASAH3: n-acylsphingosine amidohydrolase 3 alkaline ceramidase; ASAH3L: n-acylsphingosine amidohydrolase 3-like; PHCA: alkaline ceramidase 3; MFF: mitochondrial fission factor; Mfn1: mitofusin 1; Mfn2: mitofusin 2; OPA1: optic atrophy 1; P53: recombinant tumor protein p53; Bcl-2: B-cell lymphoma-2; BAX: Bcl2-Associated X.

## 4. Discussion

The present study provides comprehensive clinical and experimental evidence that n-3 polyunsaturated fatty acids (PUFAs) derived from distinct marine sources-namely fish oil (triglyceride form) and krill oil (phospholipid form)-ameliorate post-infarction cardiac dysfunction through modulation of the adiponectin–ceramide metabolic axis, with no discernible form-specific difference in therapeutic efficacy. A case-control analysis revealed that serum concentrations of specific long-chain ceramide species were markedly elevated in patients with acute myocardial infarction and exhibited a significant inverse correlation with left ventricular ejection fraction. Translation of these clinical observations to a murine model of myocardial infarction demonstrated that dietary supplementation with marine-derived n-3 PUFAs effectively reduced myocardial infarct size and preserved cardiac function. Mechanistically, n-3 PUFA intervention upregulated myocardial adiponectin expression and receptor signaling, thereby suppressing de novo ceramide synthesis via downregulation of serine palmitoyltransferase subunits SPTLC2 and SPTLC3, while concurrently promoting ceramide catabolism through ASAH1. This coordinated attenuation of pathologic ceramide accumulation restored mitochondrial dynamics, mitigated p53 and caspase-3 dependent apoptosis, and ultimately conferred improved survival and functional recovery following ischemic injury. Collectively, these findings identify the adiponectin–ceramide cascade as a critical lipid-sensing pathway underlying the cardioprotective actions of n-3 PUFAs in the post-infarction setting.

The case-control study initially established that ceramides serve as biomarkers for myocardial infarction (MI). Specifically, left ventricular ejection fraction (LVEF) exhibited a significant inverse correlation with serum ceramide concentrations, while the traditional MI biomarker troponin showed a positive correlation with ceramides—collectively indicating a link between ceramide metabolism and MI pathogenesis. Compared to healthy controls without a history of cardiovascular disease, patients with MI displayed significantly elevated levels of target ceramides, including C16:0-, C18:0-, C20:0-, and C24:1-. This observation aligns with findings from prior investigations.^[34,35]^.

Previous studies have demonstrated that n-3 PUFAs significantly reduce serum ceramide levels^[17]^. To investigate whether n-3 PUFAs from distinct sources improve post-MI cardiac function by targeting the ceramide metabolic pathway, we established a mouse MI model and systematically compared the effects of two marine-derived n-3 PUFAs—fish and krill oil—on post-MI cardiac function, mitochondrial homeostasis, and functional outcomes. The findings demonstrated that both n-3 PUFAs exerted favorable effects on infarct size reduction, ejection fraction improvement, and post-MI prognosis; however, no statistically significant differences were observed between the two interventions. This suggests that their cardioprotective effects are primarily attributed to their shared bioactive components, EPA and DHA, rather than structural differences in their lipid carriers. The structural and functional diversity of ceramides is determined by the length of their N-acyl chains. Among these, C16:0- and C18:0-have been demonstrated to fatty acid oxidation, and energy expenditure^[36–38]^. Furthermore, pathological ceramide accumulation induces apoptosis by dysregulating mitochondrial fission and fusion^[39]^. The concentrations of C16:0-, C18:0-, and total ceramides were significantly reduced in the hearts of krill oil– and fish oil–treated animals. Given that n-3 PUFAs elevate serum adiponectin levels^[40,41]^, and adiponectin treatment enhances ceramide catabolism and promotes cell survival^[42]^, we further investigated whether n-3 PUFA administration modulates cardiac adiponectin expression and thereby suppresses cardiomyocyte apoptosis.

Mechanistically, administration of n-3 PUFAs activated the adiponectin-ceramidase signaling pathway, which suppressed de novo ceramide synthesis and accelerated ceramide catabolism, thereby inhibiting cardiomyocyte apoptosis and improving myocardial function. Adiponectin is exclusively secreted by adipocytes and exerts well-characterized anti-inflammatory, anti-apoptotic, and cardioprotective effects^[43]^. Upon binding to its receptor AdipoR2, adiponectin coordinately suppresses de novo ceramide synthesis—via downregulation of the expression and activity of serine palmitoyltransferase subunits SPTLC2/3^[44,45]^—and activates ASAH1 to promote ceramide catabolism^[42,46]^. In the present study, krill oil and fish oil interventions significantly elevated cardiac protein levels of adiponectin and ASAH1, while concurrently reducing SPTLC2/SPTLC3 protein expression. Ceramide accumulation disrupts mitochondrial homeostasis, leading to imbalanced mitochondrial dynamics marked by excessive fission and impaired fusion^[47–49]^. Consistently, elevated ceramide concentrations upregulate the expression of MFF^[50,51]^; notably, both krill oil and fish oil treatments significantly attenuated MFF overexpression relative to the MI model group. Excessive mitochondrial fission and fusion defects are mechanistically linked to cardiomyocyte apoptosis and contractile dysfunction^[52,53]^. Furthermore, ceramide-driven mitochondrial dysfunction reprograms cardiomyocyte metabolism and further amplifies apoptotic signaling^[54,55]^. Collectively, this study demonstrates that krill oil and fish oil mitigate cardiomyocyte apoptosis by coordinately upregulating anti-apoptotic proteins (e.g., caspase-3) and inhibiting the activation of pro-apoptotic effectors (e.g., cleaved caspase-3). The core mechanism involves activation of the adiponectin-ceramide axis, which restores mitochondrial dynamics balance, suppresses pathological apoptosis, and ultimately improves cardiac remodeling and functional recovery following myocardial infarction (Figure 10).

**Figure 10.**
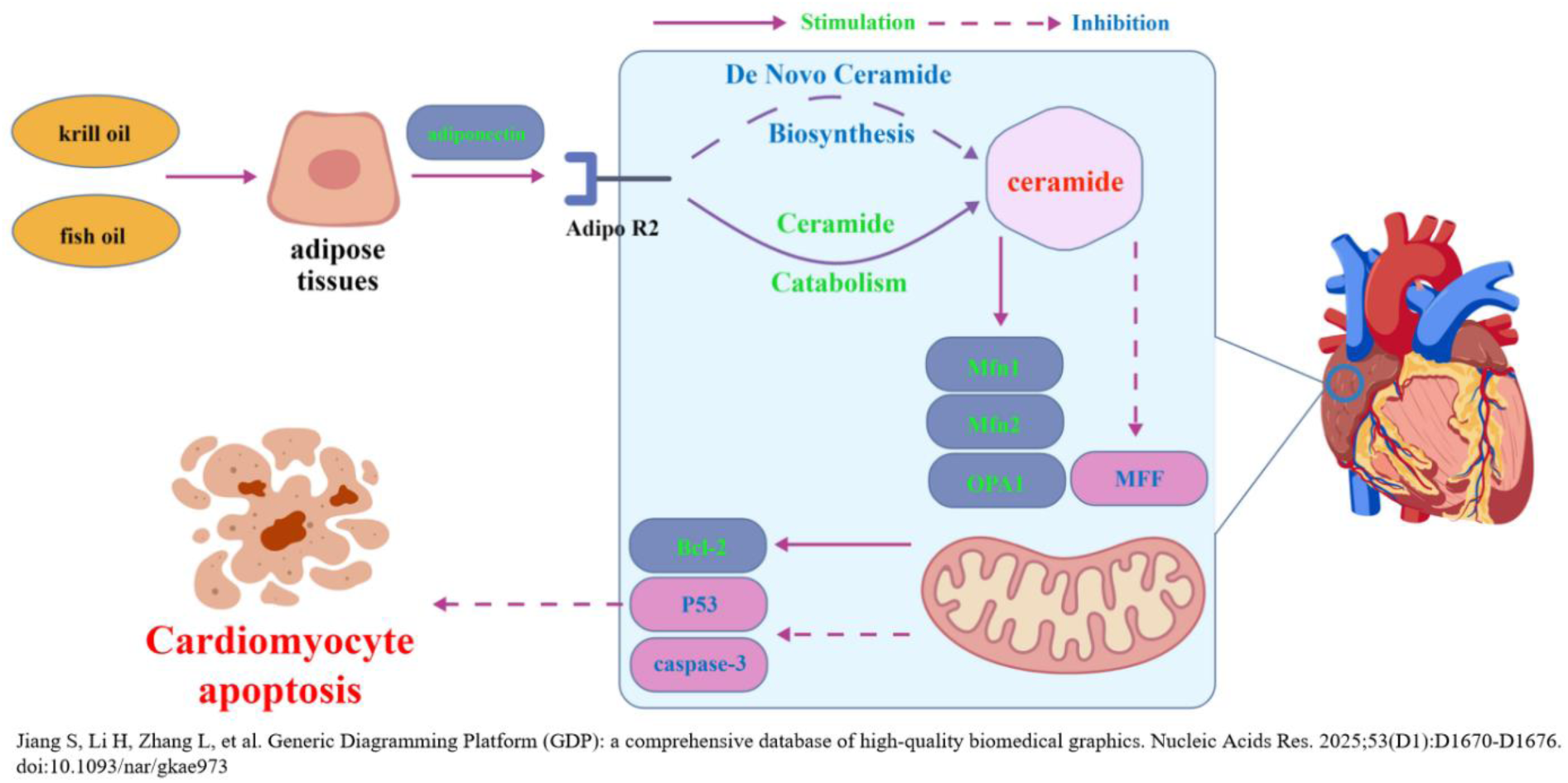
The underlying mechanism through which n-3 PUFA improves cardiac dysfunction through the adiponectin/ ceramide/ apoptosis pathway.

## 5. Limitation

This study employed a case-control design, which inherently limits the establishment of causal relationships. Additionally, the lack of long-term follow-up renders the long-term prognosis of patients uncertain. In animal experiments, although n-3 PUFA intervention was observed to modulate the adiponectin/ceramide axis and improve cardiac function, targeted inhibition of this axis via genetic or pharmacological approaches has not been performed. Thus, the necessity of the adiponectin/ceramide signaling pathway in the cardioprotective effects of n-3 PUFAs has not been directly verified—a critical gap in mechanistic validation that constitutes a major limitation of the current study.

## 6. Conclusion

This study demonstrates that n-3 PUFAs derived from fish oil and krill oil significantly improve post-MI cardiac function recovery and histopathological outcomes in C57BL/6J mice. The core mechanism involves upregulation of adiponectin expression, which sequentially inhibits SPTLC2/3—a key enzyme in the de novo ceramide synthesis pathway—and activates ASAH1. These effects synergistically reduce myocardial levels of pathogenic long-chain ceramides, ultimately restoring the dynamic balance of mitochondrial fission and fusion. This finding not only elucidates a novel lipid metabolic regulatory axis underlying the cardioprotective effects of n-3 PUFAs but also provides translational evidence for targeting ceramide metabolism to mitigate adverse post-MI outcomes.

## Acknowledgments

The authors gratefully acknowledge all patients and healthy volunteers who participated in this case–control study. We also thank the staff of the Laboratory Animal Center at Qingdao University for their expert support in animal husbandry and experimental protocol implementation.

---

**Supplementary Figure 1.**
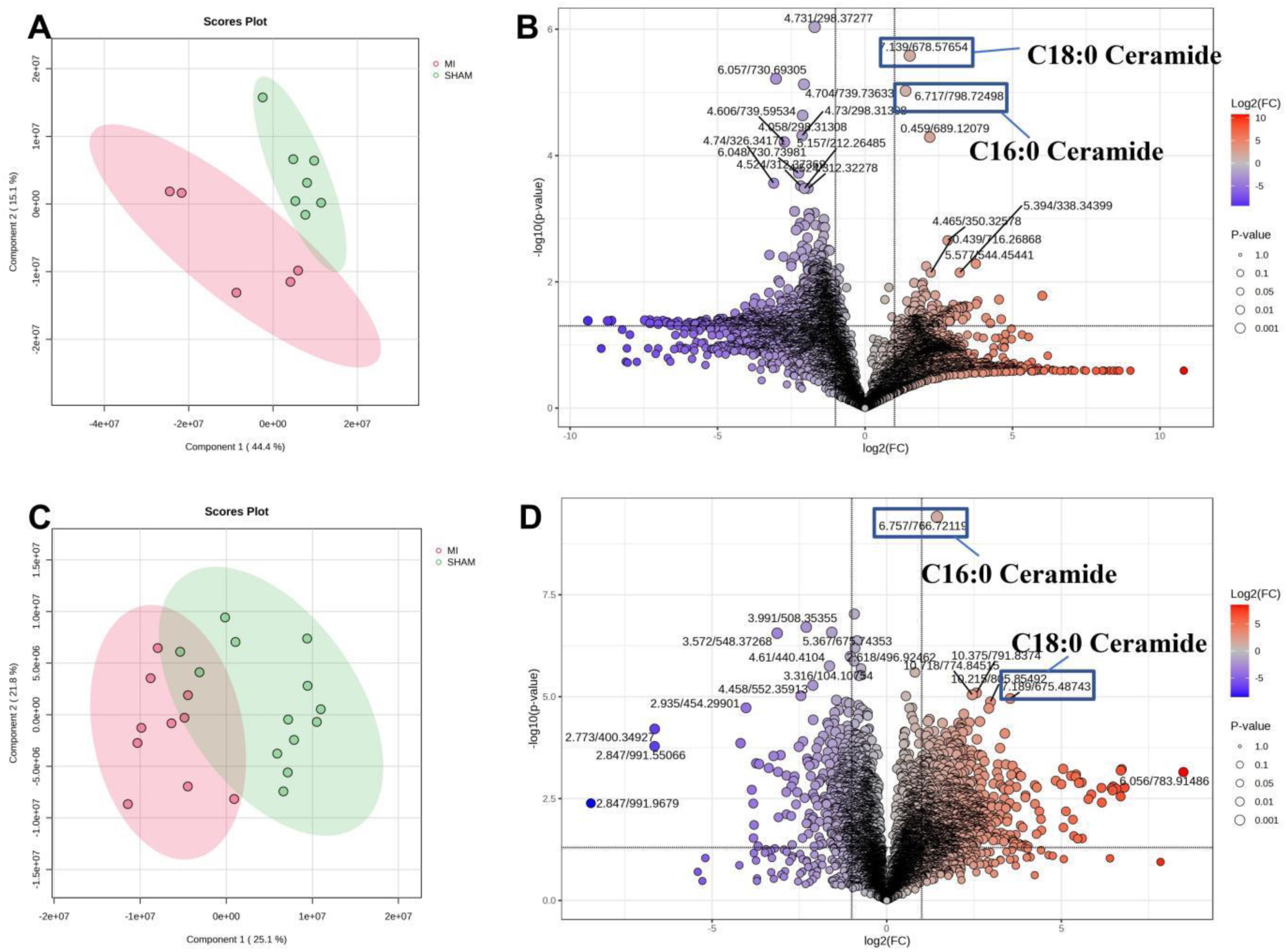
Lipidomics Analysis: (A) PCA of Cardiac Lipids in the MI Group and SHAM Group. (B) Volcano Plot Comparison of Cardiac Lipid Metabolic Data Between the MI Group and SHAM Group. (C) PCA of Serum Lipids in the MI Group and SHAM Group. (D) Volcano Plot Comparison of Serum Lipid Metabolic Data Between the MI Group and SHAM Group.

**Supplementary Table 1.** Fatty Acid Composition of the Interventions.

| Fatty acid (%) | Fish oil | Krill oil | Olive |
| --- | --- | --- | --- |
| **C16:0** | 11.75±1.20 | 18.79±1.07 | 10.51±0.12 |
| **C18:0** | 3.40±0.52 | 1.07±0.02 | 3.68±0.13 |
| **C18:1n-9** | 12.55±2.13 | 10.55±0.06 | 77.60±0.66 |
| **C18:3n-3** | 0.44±0.12 | 0.18±0.01 | -- |
| **C18:3n-6** | 0.51±0.08 | 0.62±0.02 | 0.66±0.01 |
| **C20:1n-9** | 2.15±0.05 | 0.88±0.03 | 0.26±0.01 |
| **C20:4n-6** | 2.33±0.06 | 0.65±0.04 | -- |
| **C20:5n-3** | 25.44±3.21 | 27.03±1.23 | -- |
| **C22:6n-3** | 17.16±2.25 | 11.76±1.89 | -- |

**Supplementary Table 2.** Cardiac Fatty Acid Composition.

| **Fatty acid**<br>**(%)** | **MI** | **SHAM** | **Krill oil** | **Fish oil** | ***p*-value** |
| --- | --- | --- | --- | --- | --- |
| **C4:0** | 0.058±0.062 | 0.034±0.010 | 0.014±0.008 | 0.081±0.056 | 0.051 |
| **C6:0** | 0.034±0.022<sup>ab</sup> | 0.022±0.011<sup>b</sup> | 0.015±0.013<sup>b</sup> | 0.056±0.020<sup>a</sup> | 0.026 |
| **C8:0** | 0.041±0.081 | 0.008±0.006 | 0.004±0.002 | 0.010±0.004 | 0.543 |
| **C10:0** | 0.018±0.016 | 0.016±0.010 | 0.010±0.005 | 0.014±0.004 | 0.722 |
| **C11:0** | 0.318±0.271 | 0.313±0.198 | 0.174±0.087 | 0.363±0.285 | 0.606 |
| **C13:0** | 0.576±0.234 | 0.462±0.273 | 0.552±0.502 | 0.351±0.180 | 0.777 |
| **C14:0** | 0.679±0.353<sup>b</sup> | 0.720±0.266<sup>b</sup> | 0.832±0.293<sup>b</sup> | 1.355±0.144<sup>a</sup> | 0.029 |
| **C14:1** | 0.134±0.025 | 0.138±0.044 | 0.101±0.033 | 0.112±0.008 | 0.248 |
| **C15:0** | 0.073±0.025<sup>b</sup> | 0.103±0.030<sup>b</sup> | 0.091±0.030<sup>b</sup> | 0.155±0.389<sup>a</sup> | 0.012 |
| **C15:1** | 16.562±2.350 | 14.470±2.793 | 14.696±3.981 | 13.999±0.907 | 0.524 |
| **C16:0** | 1.854±1.246 | 1.508±1.820 | 1.733±1.693 | 1.314±0.292. | 0.953 |
| **C16:1** | 0.301±0.121 | 0.301±0.077 | 0.243±0.034 | 0.186±0.082 | 0.171 |
| **C17:0** | 0.104±0.259 | 0.133±0.042 | 0.080±0.046 | 0.100±0.714 | 0.342 |
| **C17:1** | 12.129±3.847 | 12.609±3.357 | 12.241±1.522 | 10.802±1.562 | 0.660 |
| **C18:1n-9c/t** | 19.205±3.856 | 15.222±3.891 | 14.455±4.745 | 14.433±1.880 | 0.191 |
| **C18:2n-6t** | 23.304±2.423<sup>a</sup> | 23.082±1.808<sup>a</sup> | 20.039±0.394<sup>b</sup> | 20.189±1.251<sup>b</sup> | 0.015 |
| **C18:3n-6** | 0.075±0.063<sup>ab</sup> | 0.119±0.039<sup>a</sup> | 0.053±0.052<sup>b</sup> | 0.019±0.007<sup>b</sup> | 0.050 |
| **C18:3n-3** | 0.403±0.212 | 0.458±0.357 | 0.249±0.113 | 0.209±0.051 | 0.211 |
| **C20:0** | 0.257±0.079<sup>a</sup> | 0.278±0.058<sup>a</sup> | 0.184±0.045<sup>b</sup> | 0.125±0.074<sup>b</sup> | 0.019 |
| **C20:1n-9** | 0.590±0.093<sup>ab</sup> | 0.644±0.155<sup>a</sup> | 0.470±0.095<sup>b</sup> | 0.275±0.249<sup>b</sup> | 0.014 |
| **C20:2** | 0.259±0.026<sup>a</sup> | 0.244±0.033<sup>ab</sup> | 0.195±0.019<sup>b</sup> | 0.103±0.090<sup>ab</sup> | 0.016 |
| **C20:3n-6** | 0.221±0.041<sup>a</sup> | 0.234±0.041<sup>a</sup> | 0.191±0.038<sup>ab</sup> | 0.128±0.074<sup>b</sup> | 0.033 |
| **C21:0** | 0.299±0.062 | 0.299±0.039 | 0.242±0.076 | 0.181±0.088 | 0.071 |
| **C20:4n-6**<br>**(ARA)** | 7.019±1.159<sup>a</sup> | 8.041±1.924<sup>a</sup> | 3.367±0.285<sup>b</sup> | 3.538±0.517<sup>b</sup> | <0.001 |
| **C20:3n-3** | 0.398±0.423 | 0.628±0.958 | 0.678±0.641 | 0.313±0.279 | 0.815 |
| **C20:5n-3**<br>**(EPA)** | 0.091±0.065<sup>b</sup> | 0.139±0.087<sup>b</sup> | 0.546±0.232<sup>ab</sup> | 0.690±0.122<sup>a</sup> | <0.001 |
| **C22:0** | 0.109±0.059 | 0.129±0.043 | 0.084±0.029 | 0.138±0.082 | 0.474 |
| **C22:1n-9** | 0.116±0.081 | 0.115±0.052 | 0.068±0.016 | 0.146±0.106 | 0.437 |
| **C22:2** | 0.038±0.015<sup>b</sup> | 0.062±0.053<sup>b</sup> | 0.045±0.025<sup>b</sup> | 0.126±0.002<sup>a</sup> | 0.009 |
| **C23:0** | 0.034±0.024<sup>b</sup> | 0.055±0.028<sup>b</sup> | 0.057±0.030<sup>b</sup> | 0.120±0.028<sup>a</sup> | 0.004 |
| **C24:0** | 0.629±0.237<sup>b</sup> | 0.837±0.159<sup>a</sup> | 1.615±0.646<sup>a</sup> | 1.254±0.206<sup>a</sup> | 0.020 |
| **C22:6n-3**<br>**(DHA)** | 13.853±2.981<sup>b</sup> | 18.333±6.694<sup>ab</sup> | 26.161±10.180<sup>a</sup> | 27.633±2.976<sup>a</sup> | 0.018 |
| **C24:1** | 0.220±0.110<sup>b</sup> | 0.222±0.037<sup>b</sup> | 0.513±0.411<sup>b</sup> | 1.491±1.310<sup>a</sup> | 0.018 |

Effect of krill and fish oil intervention on cardiac fatty acid composition in C57BL/6J mice.
Values were presented as mean ± SD (n = 20 animals/group). Different letters indicated significant differences between groups.

## Reference

[1] Byrne RA, Rossello X, Coughlan JJ, Barbato E, Berry C, Chieffo A, Claeys MJ, Dan GA, Dweck MR, Galbraith M, Gilard M, Hinterbuchner L, Jankowska EA, Jüni P, Kimura T, Kunadian V, Leosdottir M, Lorusso R, Pedretti RFE, Rigopoulos AG, Rubini Gimenez M, Thiele H, Vranckx P, Wassmann S, Wenger NK, Ibanez B; ESC Scientific Document Group. 2023 ESC Guidelines for the management of acute coronary syndromes. Eur Heart J. 2023 Oct 12;44(38):3720–3826. doi: 10.1093/eurheartj/ehad191.

[2] Thygesen K, Alpert JS, Jaffe AS, Chaitman BR, Bax JJ, Morrow DA, et al. Fourth universal definition of myocardial infarction (2018). Eur Heart J 2019;40:237–269. 10.1093/eurheartj/ehy462.

[3] Spencer FA, Meyer TE, Gore JM and Goldberg RJ. Heterogeneity in the management and outcomes of patients with acute myocardial infarction complicated by heart failure: the National Registry of Myocardial Infarction. Circulation. 2002;105:2605–2610.

[4] Hadas Y, Katz MG, Bridges CR and Zangi L. Modified mRNA as a therapeutic tool to induce cardiac regeneration in ischemic heart disease. Wiley Interdiscip Rev Syst Biol Med. 2017;9: wsbm.1367.

[5] Lin Z and Pu WT. Strategies for cardiac regeneration and repair. Sci Transl Med. 2014;6:239rv1.

[6] Correction to: 2018 AHA/ ACC/ AACVPR/ AAPA/ ABC/ ACPM /ADA/ AGS/ APhA/ ASPC/ NLA/ PCNA Guideline on the Management of Blood Cholesterol: A Report of the American College of Cardiology/American Heart Association Task Force on Clinical Practice Guidelines. Circulation. 2023 Aug 15;148(7):e5. doi: 10.1161/CIR.0000000000001172.

[7] Mach F, Baigent C, Catapano AL, et al; ESC Scientific Document Group. 2019 ESC/EAS Guidelines for the management of dyslipidaemias: lipid modification to reduce cardiovascular risk. Eur Heart J. 2020 Jan 1;41(1):111–188. doi: 10.1093/eurheartj/ehz455.

[8] Mason RP, Libby P, Bhatt DL. Emerging mechanisms of cardiovascular protection for the omega-3 fatty acid eicosapentaenoic acid[J]. Arterioscler Thromb Vasc Biol, 2020, 40(5): 1135–1147. DOI: 10.1161/ATVBAHA.119.313286.

[9] Innes JK, Calder PC. Marine omega-3 (n-3) fatty acids for cardiovascular health: an update for 2020[J]. Int J Mol Sci, 2020, 21(4): 1362. DOI:10.3390/ijms21041362.

[10] Dixon DL. Catch of the day: icosapent ethyl for reducing cardiovascular risk[J]. Am J Med, 2020, 133(7): 802–804. DOI: 10.1016/j.amjmed.2020.03.006.

[11] Skulas-Ray AC, Wilson PWF, Harris WS, et al. Omega-3 fatty acids for the management of hypertriglyceridemia: a science advisory from the American Heart Association[J]. Circulation, 2019, 140(12): e673–e691. DOI: 10.1161/CIR.0000000000000709.

[12] Satoh N, Shimatsu A, Kotani K, et al. Purified eicosapentaenoic acid reduces small dense LDL, remnant lipoprotein particles, and C-reactive protein in metabolic syndrome[J]. Diabetes Care, 2007, 30(1): 144–146. DOI: 10.2337/dc06-1179.

[13] Decandia D, Landolfo E, Sacchetti S, Gelfo F, Petrosini L, Cutuli D. n-3 PUFA Improve Emotion and Cognition during Menopause: A Systematic Review. Nutrients. 2022 May 9;14(9):1982. doi: 10.3390/nu14091982.

[14] Kim MG, Yang I, Lee HS, Lee JY, Kim K. Lipid-modifying effects of krill oil vs fish oil: a network meta-analysis. Nutr Rev. 2020 Sep 1;78(9):699–708. doi: 10.1093/nutrit/nuz102.

[15] Ogata S, Manson JE, Kang JH, Buring JE, Lee IM, Nishimura K, Sakata Y, Danik JS, D’Agostino D, Mora S, Albert CM, Cook NR. Marine n-3 Fatty Acids and Prevention of Cardiovascular Disease: A Novel Analysis of the VITAL Trial Using Win Ratio and Hierarchical Composite Outcomes. Nutrients. 2023 Sep 30;15(19):4235. doi: 10.3390/nu15194235.

[16] Lázaro I, Rueda F, Cediel G, Ortega E, García-García C, Sala-Vila A, Bayés-Genís A. Circulating Omega-3 Fatty Acids and Incident Adverse Events in Patients With Acute Myocardial Infarction. J Am Coll Cardiol. 2020 Nov 3;76(18):2089–2097. doi: 10.1016/j.jacc.2020.08.073

[17] Chen YF, Fan ZK, Wang YP, Liu P, Guo XF, Li D. Docosahexaenoic Acid Modulates Nonalcoholic Fatty Liver Disease by Suppressing Endocannabinoid System. Mol Nutr Food Res. 2024 Apr;68(7):e2300616. doi: 10.1002/mnfr.202300616

[18] Holland W L, Brozinick J T, Wang L P, et al. Inhibition of ceramide synthesis ameliorates glucocorticoid-, saturated-fat-, and obesity-induced insulin resistance Cell Metab, 2007, 5(3): 167–79.

[19] Summers S A, Garza L A, Zhou H L, et al. Regulation of insulin-stimulated glucose transporter GLUT4 translocation and Akt kinase activity by ceramide [J]. Mol Cell Biol, 1998, 18(9): 5457–64.

[20] Turpin S M, Nicholls H T, Willmes D M, et al. Obesity-Induced CerS6-Dependent C16:0< Ceramide Production Promotes Weight Gain and Glucose Intolerance [J]. Cell Metab, 2014, 20(4): 678–86.

[21] Chaurasia B, Summers S A. Ceramides - Lipotoxic Inducers of Metabolic Disorders [J]. Trends Endocrinol Metab, 2015, 26(10): 538–50.

[22] SenthilKumar G, Zirgibel Z, Cohen KE, Katunaric B, Jobe AM, Shult CG, Limpert RH, Freed JK. Ying and Yang of Ceramide in the Vascular Endothelium. Arterioscler Thromb Vasc Biol. 2024 Aug;44(8):1725–1736. doi: 10.1161/ATVBAHA.124.321158. Epub 2024 Jun 20.

[23] Akhiyat N, Vasile V, Ahmad A, Sara JD, Nardi V, Lerman LO, Jaffe A, Lerman A. Plasma ceramide levels are elevated in patients with early coronary atherosclerosis and endothelial dysfunction. J Am Heart Assoc. 2022;11:e022852. doi: 10.1161/JAHA.121.022852.

[24] Freed JK, Durand MJ, Hoffmann BR, Densmore JC, Greene AS, Gutterman DD. Mitochondria-regulated formation of endothelium-derived extracellular vesicles shifts the mediator of flow-induced vasodilation. Am J Physiol Heart Circ Physiol. 2017;312:H1096–H1104. doi: 10.1152/ajpheart.00680.2016.

[25] De Palma C, Meacci E, Perrotta C, Bruni P, Clementi E. Endothelial nitric oxide synthase activation by tumor necrosis factor alpha through neutral sphingomyelinase 2, sphingosine kinase 1, and sphingosine 1 phosphate receptors: a novel pathway relevant to the pathophysiology of endothelium. Arterioscler Thromb Vasc Biol. 2006;26:99–105. doi: 10.1161/01.ATV.0000194074.59584.42.

[26] Teragawa H, Ueda K, Matsuda K, Kimura M, Higashi Y, Oshima T, Yoshizumi M, Chayama K. Relationship between endothelial function in the coronary and brachial arteries. Clin Cardiol. 2005;28:460–466. doi: 10.1002/clc.4960281004.

[27] R.L. Castillo, C. Arias, J.G. Farías, Omega 3 chronic supplementation attenuates myocardial ischaemia-reperfusion injury through reinforcement of antioxidant defense system in rats, Cell Biochem. Funct. 32 (2014) 274–281. 10.1002/cbf.3012.

[28] D. Richard, F. Oszust, C. Guillaume, H. Millart, D. Laurent-Maquin, C. Brou, P. Bausero, F. Visioli, Infusion of docosahexaenoic acid protects against myocardial infarction, Prostaglandins Leukot. Essent. Fat. Acids 90 (2014) 139–143. 10.1016/j.plefa.2014.01.001.

[29] Roy J, Fauconnier J, Oger C, Farah C, Angebault-Prouteau C, Thireau J, Bideaux P, Scheuermann V, Bultel-Poncé V, Demion M, Galano JM, Durand T, Lee JC, Le Guennec JY. Non-enzymatic oxidized metabolite of DHA, 4(RS)-4-F4t-neuroprostane protects the heart against reperfusion injury. Free Radic Biol Med. 2017 Jan;102:229–239. doi: 10.1016/j.freeradbiomed.2016.12.005.

[30] Shi Y, Li H, Wu T, Wang Q, Zhu Q, Guan X, Wu R. Docosahexaenoic Acid-Enhanced Autophagic Flux Improves Cardiac Dysfunction after Myocardial Infarction by Targeting the AMPK/mTOR Signaling Pathway. Oxid Med Cell Longev. 2022 Feb 27;2022:1509421. doi: 10.1155/2022/1509421.

[31] Rao SV, O’Donoghue ML, Ruel M, Rab T, Tamis-Holland JE, Alexander JH, Baber U, Baker H, Cohen MG, Cruz-Ruiz M, Davis LL, de Lemos JA, DeWald TA, Elgendy IY, Feldman DN, Goyal A, Isiadinso I, Menon V, Morrow DA, Mukherjee D, Platz E, Promes SB, Sandner S, Sandoval Y, Schunder R, Shah B, Stopyra JP, Talbot AW, Taub PR, Williams MS. 2025 ACC/AHA/ACEP/NAEMSP/SCAI Guideline for the Management of Patients With Acute Coronary Syndromes: A Report of the American College of Cardiology/American Heart Association Joint Committee on Clinical Practice Guidelines. Circulation. 2025 Apr;151(13):e771–e862. doi: 10.1161/CIR.0000000000001309.

[32] Gao E, Lei YH, Shang X, Huang ZM, Zuo L, Boucher M, Fan Q, Chuprun JK, Ma XL, Koch WJ. A novel and efficient model of coronary artery ligation and myocardial infarction in the mouse. Circ Res. 2010 Dec 10;107(12):1445–53. doi: 10.1161/CIRCRESAHA.110.223925

[33] Shahidi F, Ambigaipalan P. Omega-3 Polyunsaturated Fatty Acids and Their Health Benefits [M]. 2018 Mar: 345–81.

[34] Chenchen Tu, Lan Xie, Zhenjie Wang, et al. Association between ceramides and coronary artery stenosis in patients with coronary artery disease. Lipids Health Dis. 2020 Jun 25; 19(1): 151. doi: 10.1186s12944-020-01329-0.

[35] Huiqing Liang, Fangjiang Li, Liang Zhang, et al. Ceramides and pro-inflammatory cytokines for the prediction of acute coronary syndrome: a multi-marker approach. BMC Cardiovascular Disorders. 2024 Jan 13; 24(1): 47. doi: 10.1186/s12872-023-03690-1.

[36] Perreault L, Newsom SA, Strauss A, Kerege A, Kahn DE, Harrison KA, Snell-Bergeon JK, Nemkov T, D’Alessandro A, Jackman MR, MacLean PS, Bergman BC. Intracellular localization of diacylglycerols and sphingolipids influences insulin sensitivity and mitochondrial function in human skeletal muscle. JCI Insight. 2018 Feb 8;3(3):e96805. doi: 10.1172/jci.insight.96805.

[37] Chen TC, Benjamin DI, Kuo T, Lee RA, Li ML, Mar DJ, Costello DE, Nomura DK, Wang JC. The glucocorticoid-Angptl4-ceramide axis induces insulin resistance through PP2A and PKCζ. Sci Signal. 2017 Jul 25;10(489):eaai7905. doi: 10.1126/scisignal.aai7905.

[38] Turpin-Nolan SM, Hammerschmidt P, Chen W, Jais A, Timper K, Awazawa M, Brodesser S, Brüning JC. CerS1-Derived C18:0 Ceramide in Skeletal Muscle Promotes Obesity-Induced Insulin Resistance. Cell Rep. 2019 Jan 2;26(1):1–10.e7. doi: 10.1016/j.celrep.2018.12.031.

[39] Liu LK, Choudhary V, Toulmay A, Prinz WA. An inducible ER-Golgi tether facilitates ceramide transport to alleviate lipotoxicity. J Cell Biol. 2017 Jan 2;216(1):131–147. doi: 10.1083/jcb.201606059.

[40] Farimani AR, Hariri M, Azimi-Nezhad M, Borji A, Zarei S, Hooshmand E. The effect of n-3 PUFAs on circulating adiponectin and leptin in patients with type 2 diabetes mellitus: a systematic review and meta-analysis of randomized controlled trials. Acta Diabetol. 2018 Jul;55(7):641–652. doi: 10.1007/s00592-018-1110-6.

[41] von Frankenberg AD, Silva FM, de Almeida JC, Piccoli V, do Nascimento FV, Sost MM, Leitão CB, Remonti LL, Umpierre D, Reis AF, Canani LH, de Azevedo MJ, Gerchman F. Effect of dietary lipids on circulating adiponectin: a systematic review with meta-analysis of randomised controlled trials. Br J Nutr. 2014 Oct 28;112(8):1235–50. doi: 10.1017/S0007114514002013.

[42] Yamauchi T, Nio Y, Maki T, Kobayashi M, Takazawa T, Iwabu M, Okada-Iwabu M, Kawamoto S, Kubota N, Kubota T, Ito Y, Kamon J, Tsuchida A, Kumagai K, Kozono H, Hada Y, Ogata H, Tokuyama K, Tsunoda M, Ide T, Murakami K, Awazawa M, Takamoto I, Froguel P, Hara K, Tobe K, Nagai R, Ueki K, Kadowaki T. Targeted disruption of AdipoR1 and AdipoR2 causes abrogation of adiponectin binding and metabolic actions. Nat Med. 2007 Mar;13(3):332–9. doi: 10.1038/nm1557.

[43] Nawrocki AR, Rajala MW, Tomas E, Pajvani UB, Saha AK, Trumbauer ME, Pang Z, Chen AS, Ruderman NB, Chen H, Rossetti L, Scherer PE. Mice lacking adiponectin show decreased hepatic insulin sensitivity and reduced responsiveness to peroxisome proliferator-activated receptor gamma agonists. J Biol Chem. 2006 Feb 3;281(5):2654–60. doi: 10.1074/jbc.M505311200.

[44] Hoffman M, Palioura D, Kyriazis ID, Cimini M, Badolia R, Rajan S, Gao E, Nikolaidis N, Schulze PC, Goldberg IJ, Kishore R, Yang VW, Bannister TD, Bialkowska AB, Selzman CH, Drakos SG, Drosatos K. Cardiomyocyte Krüppel-Like Factor 5 Promotes De Novo Ceramide Biosynthesis and Contributes to Eccentric Remodeling in Ischemic Cardiomyopathy. Circulation. 2021 Mar 16;143(11):1139–1156. doi: 10.1161/CIRCULATIONAHA.120.047420.

[45] Ji R, Akashi H, Drosatos K, Liao X, Jiang H, Kennel PJ, Brunjes DL, Castillero E, Zhang X, Deng LY, Homma S, George IJ, Takayama H, Naka Y, Goldberg IJ, Schulze PC. Increased de novo ceramide synthesis and accumulation in failing myocardium. JCI Insight. 2017 Jul 20;2(14):e96203. doi: 10.1172/jci.insight.96203.

[46] Iwabu M, Yamauchi T, Okada-Iwabu M, Sato K, Nakagawa T, Funata M, Yamaguchi M, Namiki S, Nakayama R, Tabata M, Ogata H, Kubota N, Takamoto I, Hayashi YK, Yamauchi N, Waki H, Fukayama M, Nishino I, Tokuyama K, Ueki K, Oike Y, Ishii S, Hirose K, Shimizu T, Touhara K, Kadowaki T. Adiponectin and AdipoR1 regulate PGC-1alpha and mitochondria by Ca(2+) and AMPK/SIRT1. Nature. 2010 Apr 29;464(7293):1313–9. doi: 10.1038/nature08991.

[47] Kogot-Levin A, Saada A. Ceramide and the mitochondrial respiratory chain. Biochimie. 2014 May;100:88–94. doi: 10.1016/j.biochi.2013.07.027.

[48] Gao X, Lee K, Reid MA, Sanderson SM, Qiu C, Li S, Liu J, Locasale JW. Serine Availability Influences Mitochondrial Dynamics and Function through Lipid Metabolism. Cell Rep. 2018 Mar 27;22(13):3507–3520. doi: 10.1016/j.celrep.2018.03.017.

[49] Sentelle RD, Senkal CE, Jiang W, Ponnusamy S, Gencer S, Selvam SP, Ramshesh VK, Peterson YK, Lemasters JJ, Szulc ZM, Bielawski J, Ogretmen B. Ceramide targets autophagosomes to mitochondria and induces lethal mitophagy. Nat Chem Biol. 2012 Oct;8(10):831–8. doi: 10.1038/nchembio.1059.

[50] Hammerschmidt P, Ostkotte D, Nolte H, Gerl MJ, Jais A, Brunner HL, Sprenger HG, Awazawa M, Nicholls HT, Turpin-Nolan SM, Langer T, Krüger M, Brügger B, Brüning JC. CerS6-Derived Sphingolipids Interact with Mff and Promote Mitochondrial Fragmentation in Obesity. Cell. 2019 May 30;177(6):1536–1552.e23. doi: 10.1016/j.cell.2019.05.008.

[51] Hu Q, Zhang H, Gutiérrez Cortés N, Wu D, Wang P, Zhang J, Mattison JA, Smith E, Bettcher LF, Wang M, Lakatta EG, Sheu SS, Wang W. Increased Drp1 Acetylation by Lipid Overload Induces Cardiomyocyte Death and Heart Dysfunction. Circ Res. 2020 Feb 14;126(4):456–470. doi: 10.1161/CIRCRESAHA.119.315252.

[52] Hu L, Ding M, Tang D, Gao E, Li C, Wang K, Qi B, Qiu J, Zhao H, Chang P, Fu F, Li Y. Targeting mitochondrial dynamics by regulating Mfn2 for therapeutic intervention in diabetic cardiomyopathy. Theranostics. 2019 May 31;9(13):3687–3706. doi: 10.7150/thno.33684.

[53] Ma T, Huang X, Zheng H, Huang G, Li W, Liu X, Liang J, Cao Y, Hu Y, Huang Y. SFRP2 Improves Mitochondrial Dynamics and Mitochondrial Biogenesis, Oxidative Stress, and Apoptosis in Diabetic Cardiomyopathy. Oxid Med Cell Longev. 2021 Nov 8;2021:9265016. doi: 10.1155/2021/9265016.

[54] Zhang Y, Wang Y, Xu J, Tian F, Hu S, Chen Y, Fu Z. Melatonin attenuates myocardial ischemia-reperfusion injury via improving mitochondrial fusion/mitophagy and activating the AMPK-OPA1 signaling pathways. J Pineal Res. 2019 Mar;66(2):e12542. doi: 10.1111/jpi.12542.

[55] Bugger H, Pfeil K. Mitochondrial ROS in myocardial ischemia reperfusion and remodeling. Biochim Biophys Acta Mol Basis Dis. 2020 Jul 1;1866(7):165768. doi: 10.1016/j.bbadis.2020.165768.

